# A phage communication peptide alters *Bacillus subtilis* colony development and promotes sporulation

**DOI:** 10.64898/2026.08.23.746533

**Authors:** Bat-El Hagbi-Lazar, Zoe Levi, Meital Shema-Mìzrachi, Ronit Suissa, Zohar Tik, Shaked Uzi-Gavrilov, Lara Holoidovsky, Shira Omer Bendori, Avigdor Eldar, Michael M. Meijler

**Author notes:** Correspondence: Bat-El Hagbi-Lazar; Michael M. Meijler.

## Abstract

Temperate *Bacillus* phages use arbitrium peptides to coordinate lysis–lysogeny decisions, but whether the mature communication peptide can be sensed directly by *Bacillus subtilis* and affect its physiology and behavior is unknown. Here we show that the φ3T arbitrium peptide SAIRGA elicits a sequence- and stereochemistry-dependent response in *Bacillus subtilis* that is strongly expressed in surface-grown colony biofilms but is not accompanied by comparable changes in planktonic growth or static-liquid pellicle morphology. The response persists in the absence of AimR, the canonical arbitrium receptor. Within colonies, SAIRGA alters spatial *PtapA* activity and increases heat-resistant spore formation without increasing total viable cell yield. Untargeted metabolomics reveals broad dose-dependent remodeling that tracks peptide activity, while program-level proteomics independently converges on late-sporulation and mature-spore-associated states. This study highlights how a phage-derived peptide may act as a signal, enabling the host to pivot toward a survival-focused developmental state.

## Main

Chemical communication allows microbial populations to coordinate collective behaviours and developmental decisions^1–3^. In *Bacillus subtilis*, extracellular peptide signalling is deeply integrated into this regulatory landscape. Peptide-responsive systems including ComQXPA and Rap–Phr^4,5^ contribute to the control of competence, motility, matrix production, specialized metabolism and sporulation, allowing information from the extracellular environment and surrounding population to influence cell fate^6–10^. These processes generate phenotypically heterogeneous communities rather than uniform population-wide responses, making *B. subtilis* a useful model for studying how extracellular chemical information is translated into developmental states.

Bacteriophages also use chemical communication to coordinate collective decisions. The arbitrium system, first identified in temperate *Bacillus*-infecting SPβ-group phages, uses a secreted peptide signal to convey information about previous infections and regulate the choice between lytic growth and lysogeny^11,12^. In phage φ3T, *aimP* encodes the precursor of the mature hexapeptide SAIRGA. The peptide accumulates during successive infections, is imported into bacterial cells, and binds the phage-encoded intracellular receptor AimR, thereby altering the AimR–AimX regulatory circuit and favoring lysogeny as the signal accumulates^11,13,14^. Related arbitrium systems are widespread among phages and other mobile genetic elements and can also contribute to the regulation of prophage induction^12,15,16^. Structurally, AimR shares features with intracellular RRNPP-family peptide receptors, illustrating the convergent use of peptide recognition in phage and bacterial regulatory systems^13,14^.

The established role of arbitrium is therefore to transmit information within viral regulatory circuits. Yet its mature peptide is released into an extracellular environment occupied by the bacterial host, in which short peptides already act as potent physiological signals. This raises a distinct question: can an arbitrium peptide itself elicit a bacterial response outside of the canonical viral circuit? In particular, it remains unclear whether exposure to a defined mature phage communication peptide, in the absence of an experimentally imposed infection, is sufficient to alter host physiology or developmental state.

The physiological context of the receiving population may be important for such a response. *B. subtilis* colony biofilms are spatially and developmentally heterogeneous communities in which distinct cell states emerge across changing local environments^6,7^. Matrix-associated activity is not uniformly distributed through a developing colony. Matrix production can form a propagating peripheral front, followed spatially and temporally by a second front of sporulating cells, revealing coordinated progression between developmental programs within an otherwise genetically homogeneous population^17^. Colony development therefore provides a setting in which an extracellular cue could change the organization of bacterial cell states without necessarily producing a uniform effect on growth or on a single regulatory output.

Here, we asked whether the mature φ3T arbitrium peptide SAIRGA can alter *B. subtilis* physiology independently of experimentally imposed phage infection. We show that SAIRGA elicits an AimR-independent, context-dependent response in surface-grown colony biofilms that alters colony development and biases the population toward sporulation. These findings reveal that a defined phage communication peptide can influence bacterial developmental state outside its canonical viral signaling circuit.

## Results

### SAIRGA induces a sequence- and context-dependent biofilm response

To determine whether a phage communication peptide can alter *B. subtilis* colony biofilm development, we exposed NCIB 3610 colony biofilms to the φ3T arbitrium peptide SAIRGA and sequence-related control peptides. After 72 h on MSgg agar, SAIRGA produced a pronounced change in colony morphology accompanied by a concentration-dependent increase in colony area (Fig. 1a–c). Mean colony area increased from 377 ± 74 mm² in untreated controls to 1,035 ± 179 mm² at 5 µM SAIRGA and 2,500 ± 172 mm² at 20 µM SAIRGA. The response was strongly dependent on peptide sequence. GAIRGA, in which the N-terminal serine of SAIRGA is replaced by glycine, retained activity and produced a colony area comparable to SAIRGA at 20 µM (2,503 ± 133 mm²). By contrast, SAIRGQ, carrying a C-terminal Ala-to-Gln substitution, did not reproduce the phenotype and remained comparable to untreated colonies (388 ± 58 mm² at 20 µM; Fig. 1b,c, Supplementary Table 1). Thus, the tested N-terminal substitution was tolerated, whereas the tested C-terminal substitution abolished the colony phenotype. Because the pronounced colony expansion could reflect a general stimulation of bacterial growth, we next examined planktonic growth across a broad range of peptide concentrations. In both MSgg and LB, growth trajectories remained highly similar across SAIRGA concentrations through lag phase, exponential growth, and stationary phase (Fig. 1d, Extended Data Fig. 1). Similar growth profiles were observed with the sequence-control peptides GAIRGA and SAIRGQ (Extended Data Fig. 1). We next asked whether the response extended to another B. subtilis biofilm context. In static MSgg cultures, SAIRGA-treated and untreated cells formed pellicles with no obvious difference in macroscopic architecture after 72 h (Fig. 1e and Supplementary Fig. 2). Together, these observations argue against a simple stimulation of bacterial growth as the basis of colony expansion and show that the prominent macroscopic response is strongly dependent on growth context, emerging in surface-grown colony biofilms but not as a comparable phenotype in planktonic cultures or static-liquid pellicles.

**Fig. 1.**
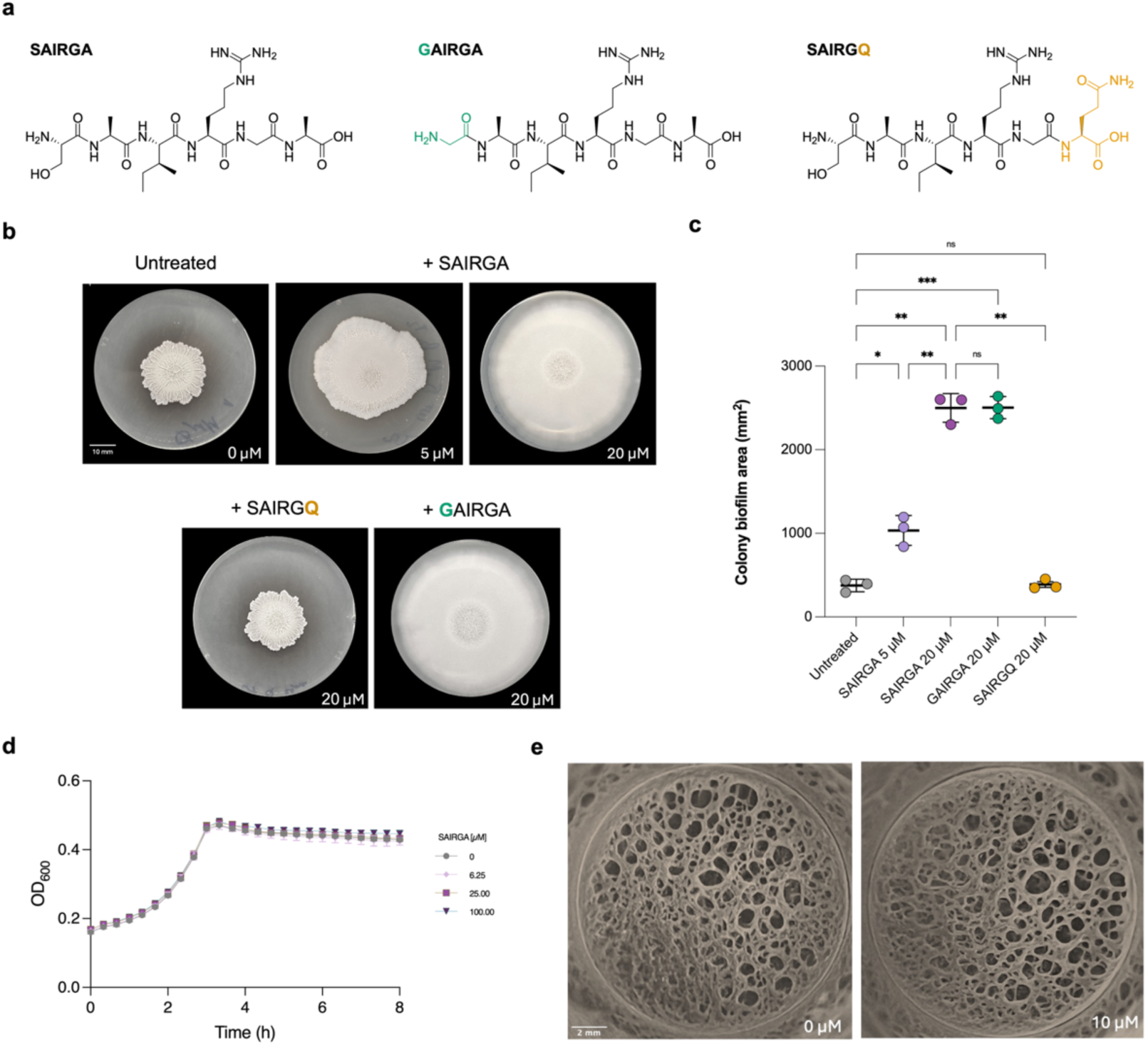
| SAIRGA induces sequence- and context-dependent remodeling of *B. subtilis* colony biofilms. **a,** Chemical structures of SAIRGA (Ser-Ala-Ile-Arg-Gly-Ala) and the sequence control peptides GAIRGA and SAIRGQ. Substitutions relative to SAIRGA are highlighted: Ser1→Gly (GAIRGA, green) and Ala6→Gln (SAIRGQ, amber). Synthetic peptide characterization is shown in Supplementary Fig. 1. **b,** Representative colony biofilms of *B. subtilis* NCIB 3610 grown on MSgg agar for 72 h at 30 °C without peptide or with SAIRGA (5 or 20 µM), GAIRGA (20 µM) or SAIRGQ (20 µM). Images are representative of three independent biological replicates; all images are shown at the same magnification. Scale bar, 10 mm. **c,** Colony biofilm area quantified from calibrated plate images. Points represent independent biological replicates (n = 3 per condition); bars show mean ± s.d. Significance was assessed by Welch’s ANOVA followed by Games–Howell multiple-comparisons testing. ns, not significant; *P < 0.05; **P < 0.01; ***P < 0.001. Additional morphometric parameters are provided in Supplementary Table 1. **d,** Planktonic growth of *B. subtilis* NCIB 3610 in liquid MSgg supplemented with the indicated concentrations of SAIRGA. OD₆₀₀ was recorded at 20-min intervals at 37 °C; the first 8 h, spanning lag phase, exponential growth and stationary phase, are shown. Data are mean ± s.d. of three independent biological replicates. The full concentration series in MSgg and LB and the corresponding GAIRGA and SAIRGQ experiments are shown in Extended Data Fig. 1. **e,** Representative top-view images of pellicles formed in static MSgg after 72 h at 30 °C without peptide or with 10 µM SAIRGA. Images are representative of three independent biological replicates; all images are shown at the same magnification. Scale bar, 2 mm. Side-view images and qualitative crystal-violet staining are shown in Supplementary Fig. 2.

### SAIRGA-induced colony remodeling depends on peptide stereochemistry but not through canonical AimR signaling

Because NCIB 3610 carries the SPβ prophage^18^, we first examined whether the SAIRGA-induced colony phenotype might be related to an endogenous arbitrium system. SPβ produces the arbitrium peptide GMPRGA^16^, which retains the C-terminal RGA motif found in SAIRGA but differs substantially at its N-terminus. GMPRGA produced a pronounced colony expansion at 20 µM, increasing mean colony area from 391 ± 79 mm² in untreated controls to 1,546 ± 302 mm² (Fig. 2a). However, the sequence-control peptide GAIRGA produced a similarly strong response (1,231 ± 85 mm²), as did SAIRGA itself (1,018 ± 172 mm²), and colony area did not differ significantly among the three active peptides. Thus, the apparently stronger response to GMPRGA was not unique to the endogenous SPβ peptide. Notably, both GMPRGA and GAIRGA carry glycine at their N-terminus, a less sterically bulky and less polar residue than the N-terminal serine of SAIRGA, raising the possibility that N-terminal sequence features contribute to response magnitude rather than the effect reflecting specific recognition of the SPβ peptide. We next examined whether peptide stereochemistry affected specific cellular uptake or recognition of the peptides. D-SAIRGA retains the amino-acid composition and sequence order of SAIRGA but is composed entirely of D-amino acids and was therefore used as a stereochemical probe to examine its effect on cellular entry or molecular recognition. Whereas L-SAIRGA increased colony area from 421 ± 45 mm² in untreated colonies to 857 ± 88 mm², D-SAIRGA did not reproduce the phenotype (513 ± 110 mm²; Fig. 2b). D-SAIRGA remained comparable to untreated colonies and was significantly less active than L-SAIRGA. This stereochemical requirement is therefore consistent with selective transport or recognition by a putative receptor and argues against a purely nonspecific extracellular effect, although this experiment alone cannot distinguish impaired uptake from altered molecular recognition. In the canonical φ3T arbitrium pathway, SAIRGA is imported into the bacterial cell and recognized by the phage-encoded receptor AimR^11,13,14^, which controls the downstream AimR–AimX regulatory circuit. We therefore asked whether the same receptor was required for the colony response. SAIRGA remained strongly active in a Δ*aimR* background, increasing mean colony area from 630 ± 42 mm² in untreated colonies to 2,513 ± 248 mm² at 20 µM (Fig. 2c). Thus, while the colony response is dependent on the stereochemistry of the signaling peptide and consistent with a specific cellular recognition process, it does not require AimR. The phenotypical change therefore cannot be explained by the known AimR-dependent arm of canonical arbitrium signalling, pointing instead to an alternative route by which *B. subtilis* responds to the phage-derived peptide.

**Fig. 2.**
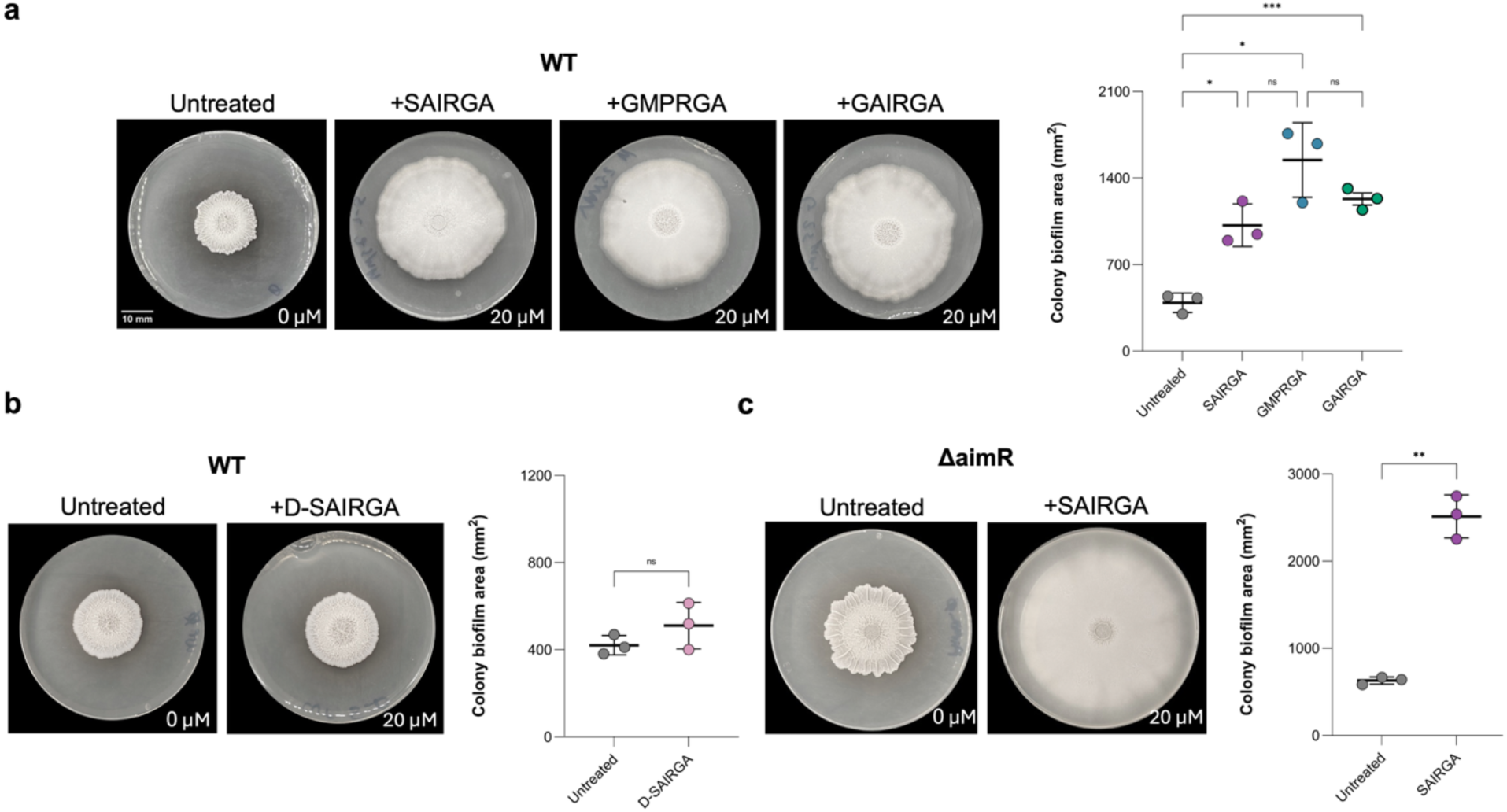
| SAIRGA-induced colony remodeling depends on peptide stereochemistry but not canonical AimR signaling. **a,** Colony biofilm area quantification (left) and representative colony biofilms (right) of wild-type *B. subtilis* NCIB 3610 grown on MSgg agar for 72 h at 30 °C without peptide or with SAIRGA, GMPRGA or GAIRGA (20 µM each). GMPRGA is the arbitrium peptide associated with the endogenous SPβ prophage, whereas GAIRGA is a sequence-control peptide carrying an N-terminal Ser1→Gly substitution relative to SAIRGA. **b,** Colony biofilm area quantification (left) and representative colony biofilms (right) of wild-type *B. subtilis* grown without peptide or with L-SAIRGA or its all-D enantiomer, D-SAIRGA (20 µM each). **c,** Colony biofilm area quantification (left) and representative colony biofilms (right) of a Δ*aimR* strain grown without peptide or with 20 µM SAIRGA. Points represent independent biological replicates (n = 3 per group); bars show mean ± s.d. Statistical significance was assessed by Welch’s ANOVA followed by Games–Howell multiple-comparisons testing for a and b, and by Welch’s *t*-test for c. ns, not significant; \**P* < 0.05; **P < 0.01; \*\*\**P* < 0.001. Scale bar, 10 mm; all colony images are shown at the same magnification. Peptide characterization is shown in Supplementary Fig. 3; additional morphometric parameters in Supplementary Table 2.

### SAIRGA shifts colony developmental state toward sporulation without increasing viable cell yield

The strong colony-specific phenotype, together with the absence of a comparable effect on planktonic growth or static-liquid pellicles, suggested that SAIRGA might alter developmental programs that emerge in surface-grown communities rather than simply affecting growth. We therefore examined two complementary aspects of colony development: the spatial organization of the matrix-associated PtapA program and the extent of sporulation, providing distinct readouts of developmental organization within the structured community. Untreated PtapA-yfp colony biofilms displayed a sharply patterned fluorescent edge, with high local fluorescence peaks around the colony perimeter (Fig. 3a–c). SAIRGA visibly altered this organization, producing a broader and more diffuse fluorescent band at the edge. A mixed-effects analysis that retained the biofilm as the biological replicate while accounting for nested spatial sampling confirmed that the quantitative effect was concentrated at the colony edge (Fig. 3d). Mean edge fluorescence increased by approximately 20% (5,117 to 6,122 arbitrary units; Šídák-adjusted P = 0.0100), whereas maximum edge fluorescence decreased by approximately 34% (15,346 to 10,093 arbitrary units; adjusted P = 0.0011). No significant treatment differences were detected in the intermediate or center zones. Radial intensity profiles were consistent with this redistribution, showing broader edge-associated fluorescence in SAIRGA-treated biofilms rather than the sharp high-intensity profile characteristic of untreated colonies (Fig. 3e). Thus, SAIRGA altered the spatial pattern of PtapA promoter activity rather than uniformly increasing or decreasing reporter expression. We next asked whether this spatial reorganization was accompanied by a change in developmental outcome. To distinguish a change in developmental state from a simple increase in cell yield, we quantified both total viable cells and heat-resistant CFU in 72-h colony biofilms. Despite the pronounced expansion of SAIRGA-treated colonies, total viable CFU per colony did not differ detectably between untreated and treated biofilms (3.67 × 10⁹ versus 4.67 × 10⁹ CFU per colony; P = 0.63; Fig. 3f). Thus, the larger colony footprint was not accompanied by a corresponding increase in viable cell number. In contrast, heat-resistant CFU increased approximately 4.8-fold, from 1.63 × 10⁸ to 7.82 × 10⁸ CFU per colony following SAIRGA treatment (P = 0.0075; Fig. 3f). Sporulation efficiency, calculated for each biological replicate as the fraction of heat-resistant CFU among total viable CFU, increased from 4.8 ± 1.7% in untreated colonies to 19.0 ± 7.1% with SAIRGA (Fig. 3f). Together, these results show that SAIRGA alters the developmental composition of the colony, increasing the representation of heat-resistant spores without increasing total viable cell yield.

**Fig. 3.**
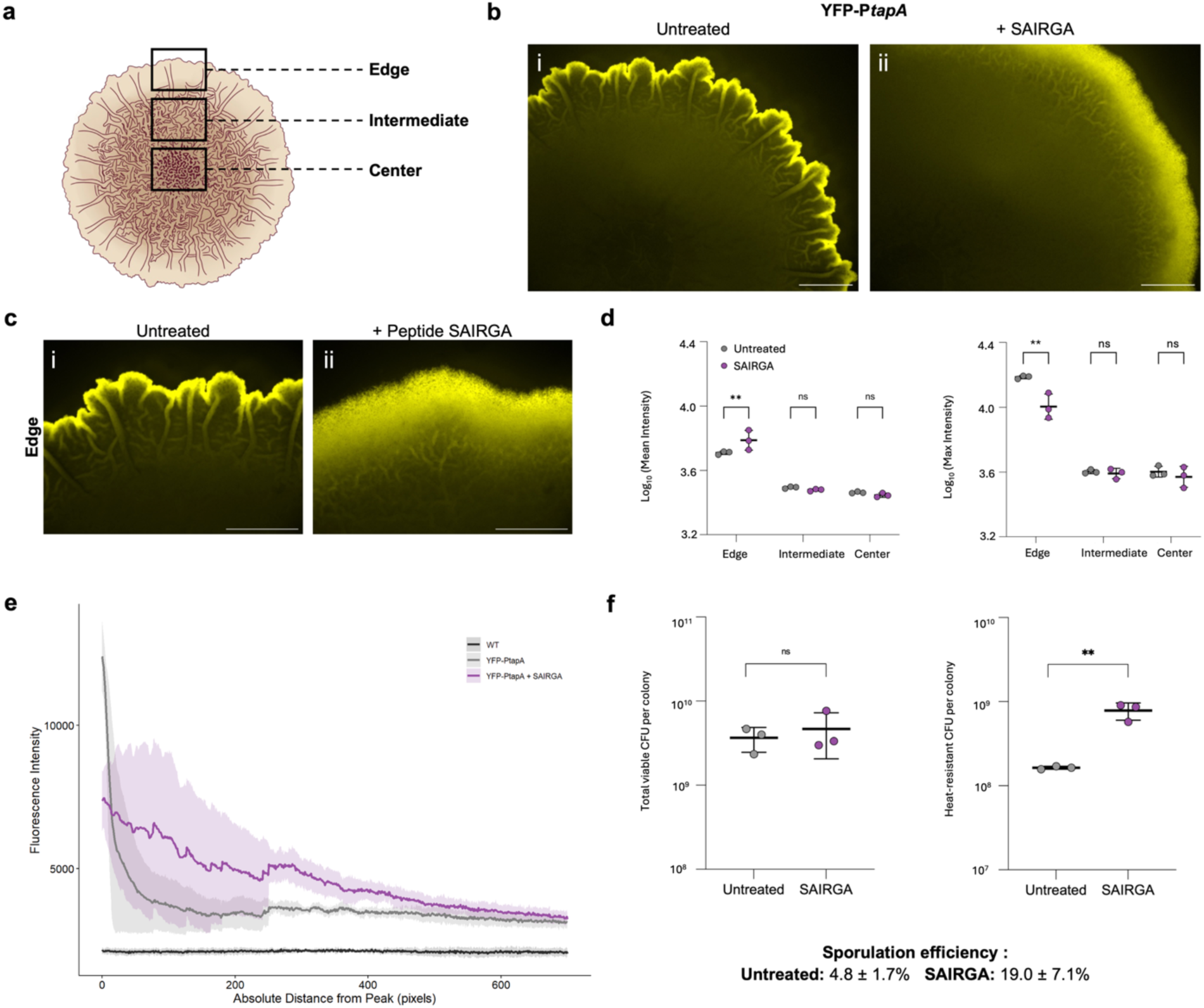
| SAIRGA alters spatial PtapA activity and promotes sporulation in colony biofilms. **a,** Schematic representation of the colony biofilm radial architecture and the edge, intermediate, and center zones used for spatial fluorescence analysis. **b,** Representative fluorescence images of PtapA-yfp colony biofilms grown without peptide or with 20 µM SAIRGA. **c,** Representative higher-magnification images of the colony edge under the same conditions. Scale bars, 2 mm. **d,** Mean (left) and maximum (right) PtapA-yfp fluorescence intensity across edge, intermediate, and center zones. Points represent three independent colony biofilms per condition after aggregation of nested spatial/technical measurements. Statistical significance was assessed using a mixed-effects analysis followed by Šídák multiple-comparisons testing of untreated versus SAIRGA-treated biofilms within each zone. ns, not significant; **P < 0.01. **e,** Radial fluorescence intensity profiles extending from the edge-associated fluorescence peak towards the colony interior. Two spatial line profiles were obtained from each of three independent biofilms per condition. Solid lines show the mean and shaded ribbons show s.d. across the six spatial profiles; profiles are displayed descriptively to 700 pixels so that all profiles contribute across the plotted range. The non-fluorescent wild-type strain is shown as a background control. **f,** Total viable CFU per 72-h colony biofilm (left) and heat-resistant CFU recovered after heat treatment at 80 °C for 10 min (right) in untreated and SAIRGA-treated samples. Points represent three independent biological replicates; each biological replicate is the mean of three technical plating replicates. Bars show mean ± s.d. CFU values were analysed after log10 transformation using unpaired t-tests with Welch’s correction. Sporulation efficiency, calculated for each biological replicate as heat-resistant CFU divided by total viable CFU × 100, increased from 4.8 ± 1.7% in untreated biofilms to 19.0 ± 7.1% with SAIRGA and is reported descriptively as a derived ratio. ns, not significant; \*\***P** < 0.01.

### SAIRGA induces dose-dependent metabolic remodeling

Because SAIRGA altered colony architecture and developmental output without increasing total viable cell yield, we next asked whether this developmental shift was accompanied by broader metabolic remodeling. To define the molecular response to SAIRGA across peptide doses, we profiled *B. subtilis* biofilm extracts by untargeted LC–HRMS/MS at SAIRGA concentrations of 1, 5, 10 and 50 µM, alongside untreated controls (Extended Data Fig. 8). Following feature detection, blank filtering and chromatographic curation, 349 high-confidence features were retained for quantitative analysis (Supplementary Table 3). PCA separated the treatment groups along two axes: PC1 (38.3%) distinguished 50 µM SAIRGA from all other conditions, while PC2 (12.8%) captured a graded separation across the intermediate concentrations (Fig. 4a). Global separation across the five conditions was significant (PERMANOVA, pseudo-*F* = 5.82, R² = 0.538, *P* = 0.001), and each SAIRGA concentration differed significantly from untreated controls (*P* = 0.009 for each pairwise comparison). The fraction of metabolomic variance associated with treatment increased across the dose series, from R² = 0.217 at 1 µM to 0.293, 0.349 and 0.573 at 5, 10 and 50 µM, respectively (Extended Data Fig. 2, Supplementary Table 4). The number of features reaching individual FDR significance also increased with concentration: no feature survived FDR correction at 1 µM, whereas 88, 103 and 193 features were significantly altered at 5, 10 and 50 µM, respectively (Welch’s *t*-test, Benjamini–Hochberg FDR < 0.05; Fig. 4b, Supplementary Table 5). This did not indicate an absence of response at the lowest dose. The metabolome was already significantly separated from untreated controls at 1 µM, and, among the 161 features subsequently identified as dose-responsive across the concentration series, 115 had already shifted in the direction of their overall trend at 1 µM. Because the response strengthened across the concentration series, we next asked which individual features changed consistently with SAIRGA dose. Spearman rank-correlation analysis across the 349-feature dataset identified 161 features with significant dose-response trends (FDR < 0.05), comprising 81 increasing and 80 decreasing features. Of these, 144 (89.4%) changed in the expected direction in at least three of the four consecutive dose steps, and 61 were strictly monotonic across all five conditions. The strongest trends and six representative strictly monotonic profiles are shown in Extended Data Fig. 3, whereas the complete 161-feature heatmap is shown in Extended Data Fig. 4 and tabulated in Supplementary Table 6.

**Fig. 4.**
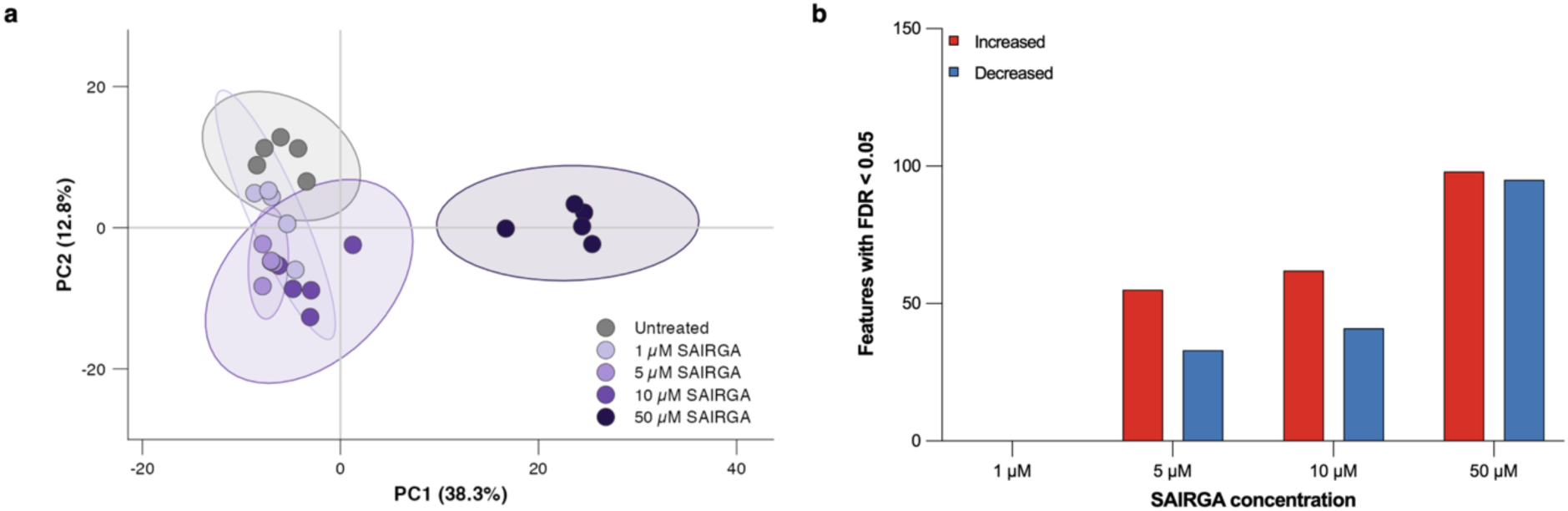
| SAIRGA induces a broad, dose-dependent shift in the *B. subtilis* colony-biofilm metabolome. **a,** Principal component analysis of 349 untargeted metabolomic features across SAIRGA concentrations (1–50 µM) and untreated controls. PC1 (38.3%) separates 50 µM SAIRGA from all other conditions, whereas PC2 (12.8%) captures a graded separation across the intermediate concentrations. Shaded regions are 95% data ellipses describing the spread of each group; statistical separation is assessed by PERMANOVA (pseudo-F = 5.82, R² = 0.54, P = 0.001 overall; P = 0.009 for each concentration versus untreated controls). Data were log₂-transformed, imputed for non-detects, and PQN- normalised. **b,** Number of features significantly altered at each concentration relative to untreated controls (Welch’s t- test with Benjamini–Hochberg correction, FDR < 0.05). Red, increased; blue, decreased. At 1 µM, no individual feature survived FDR correction, although the metabolome as a whole was already significantly separated from untreated controls (PERMANOVA P = 0.009, R² = 0.22). n = 5 independent biological replicates per condition.

### Metabolic remodeling tracks peptide activity

We next asked whether this metabolic response tracked peptide activity rather than peptide exposure alone. Using a shared feature table containing SAIRGA, the active sequence variant GAIRGA, and common untreated controls, both active peptides altered the metabolome across concentrations, although the number of significant features differed between peptides and doses. At 10 µM, fold changes across all shared features were strongly correlated between SAIRGA and GAIRGA (Pearson r = 0.863), supporting a broadly concordant metabolic response (Extended Data Fig. 5a,b). We then tested this relationship in an independent specificity dataset comparing SAIRGA with the inactive C-terminal variant SAIRGQ (Extended Data Fig. 5c). SAIRGA versus untreated controls yielded 97 features with nominal *P* < 0.05, substantially exceeding the response observed under 5,000 label permutations (empirical *P* = 0.0018). By contrast, SAIRGQ versus untreated controls yielded only two nominally significant features (empirical *P* = 0.8108). The direct SAIRGA–SAIRGQ comparison yielded 40 nominally significant features (empirical P = 0.0322), indicating that the two peptides produced distinguishable metabolomic states within the same experiment. In the unfiltered dataset, exogenously supplied SAIRGA was detected in peptide-containing agar samples across the concentration series, with increasing signal at higher nominal peptide concentrations, but in only 1 of 20 72-h colony-containing plate extracts spanning the same 1–50 µM concentration series.

### SAIRGA remodels specialized metabolism across multiple chemical families

To place the dose-responsive features in chemical context, we constructed a feature-based molecular network from Experiment 1 MS2 spectra. The resulting GNPS2 network contained 605 nodes, of which 471 formed connected molecular families and 134 were singletons (Extended Data Fig. 6a). GNPS2 spectral-library searching provided 281 library-based annotations, which were complemented by SIRIUS/CSI:FingerID structure prediction and CANOPUS chemical-class assignment using the same MS2 dataset. Additional inclusion-list acquisitions from selected stored extracts were used to obtain MS2 spectra for prioritised features and refine structural annotation; these follow-up acquisitions were used for annotation only and did not contribute quantitative abundance measurements. A complementary presence/absence analysis was evaluated in parallel to identify features dominated by condition-specific detection rather than continuous abundance; this exploratory branch was not carried forward into the biological conclusions. Nodes corresponding to the added peptides (retention time 0.47 min, annotated by CANOPUS as oligopeptides) were present in the network but were not retained in the quantitative feature table.

The dose-responsive features were not randomly distributed across the molecular network. Several connected families showed strongly directional responses (Extended Data Fig. 6b,c), including bacillaene and its derivatives (8 of 9 nodes increasing), N-acyl amino-acid lipids (15 of 18) and short peptides (4 of 4). Lysophospholipids showed a unidirectional but partial response, with 8 of 22 nodes increasing and none decreasing, whereas carbohydrate-associated features predominantly decreased. Notably, tryptophan-related chemistry showed a structured bidirectional response rather than a uniform shift. A molecular-network component containing a spectral-library match to tryptophan decreased strongly with SAIRGA dose (m/z 205.0973, ρ = −0.835, FDR = 1.61 × 10⁻⁶), together with a related indole feature (m/z 188.0707, ρ = −0.867, FDR = 2.61 × 10⁻⁷). In contrast, structurally distinct tryptophan-containing amide- and peptide-like features increased with dose, including features at m/z 431.2659 (ρ = 0.902), 488.2877 (ρ = 0.883) and 247.1079 (ρ = 0.867); SIRIUS structure prediction for the latter favored N-acetyltryptophan. These opposing trends indicate redistribution of tryptophan-derived chemistry across the SAIRGA concentration series rather than a uniform change in a single tryptophan-associated metabolite pool. The metabolomic shift resolved in Fig. 4 was therefore accompanied by structured changes within several chemically related molecular families (Supplementary Table 6).

The network also contained established *B. subtilis* specialized metabolites — plipastatin, surfactin and bacillibactin — alongside the dose-responsive bacillaene family (Extended Data Fig. 6d). Because the discovery acquisition was designed for broad metabolome coverage rather than optimal analysis of large lipopeptides, we next examined these pathways in an independently grown experiment using an extended chromatographic program and both positive- and negative-ionization measurements. SAIRGA and the inactive sequence variant SAIRGQ were tested at 20 µM alongside untreated controls in the same experiment, so that any response could be assessed against a peptide that does not induce the colony phenotype.

Targeted profiling showed a broadly upward pattern across specialized-metabolite families under SAIRGA, encompassing bacillaene, surfactin and plipastatin congeners, whereas bacillibactin was the only targeted metabolite showing a decrease relative to untreated controls (0.89-fold; Fig. 5e; Supplementary Table 7). No measurement of SAIRGQ differed significantly from untreated controls. Within this pattern, bacillaene provided the clearest response: SAIRGA increased bacillaene abundance 1.66-fold in positive ionization mode (SAIRGA versus untreated, P = 0.0042; SAIRGA versus SAIRGQ, P = 0.0084), the increase was also observed in negative ionization mode using a different internal standard (1.59-fold; P = 0.0069 and P = 0.0030, respectively; Fig. 5a,b). Plipastatin C14 likewise increased 1.31-fold relative to untreated controls (P = 0.044) and to SAIRGQ (P = 0.037; Fig. 5c). Surfactin C14 increased 1.66-fold relative to untreated controls (P = 0.020), although the direct SAIRGA–SAIRGQ comparison was not significant (P = 0.32; Fig. 5d). The broader congener series showed the same tendency towards higher lipopeptide abundance but with greater variability, and most individual measurements did not reach significance (Extended Data Fig. 7). Together, the network and targeted measurements support selective remodeling of specialized metabolism by SAIRGA, anchored by a reproducible bacillaene response and extending to representatives of the plipastatin and surfactin pathways.

**Fig. 5.**
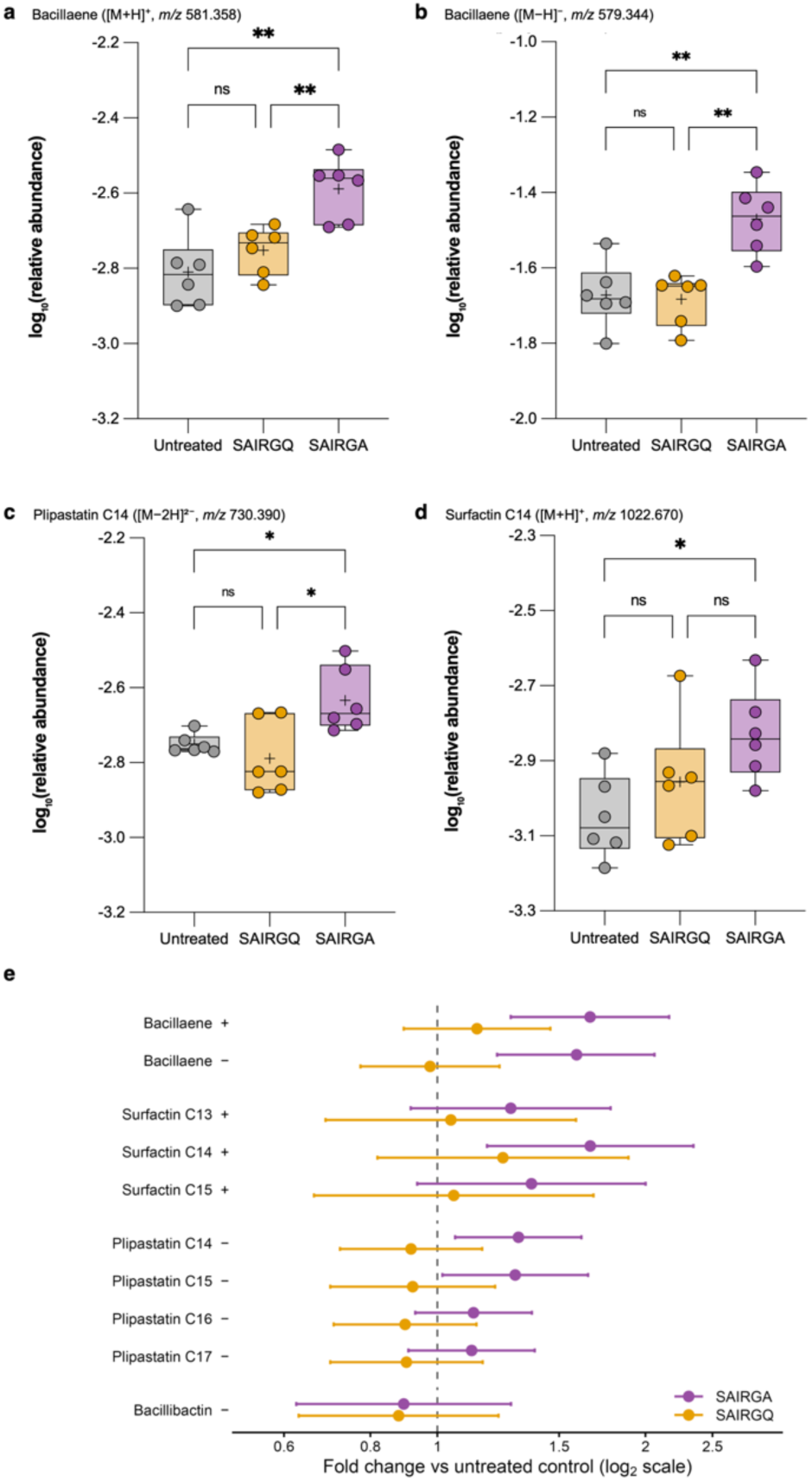
| SAIRGA increases the abundance of specialized metabolites from three biosynthetic gene clusters. Targeted LC–HRMS quantification in *B. subtilis* colony biofilms treated with SAIRGA (20 µM), SAIRGQ (20 µM) or left untreated. Positive-ionization measurements were normalized to terfenadine and negative-ionization measurements to chlorpropamide; values are log₁₀-transformed. n = 6 biological replicates per condition. **a–d,** Box plots show the median (centre line), interquartile range (box) and full data range (whiskers), with individual biological replicates overlaid; crosses indicate group means. **a,** Bacillaene ([M+H]⁺, *m/z* 581.358, ESI+). **b,** Bacillaene ([M−H]⁻, *m/z* 579.344, ESI−), providing an orthogonal measurement of the same metabolite in a second ionization mode and against a different internal standard. **c,** Plipastatin C14 ([M−2H]²⁻, *m/z* 730.390, ESI−). **d,** Surfactin C14 ([M+H]⁺, *m/z* 1022.670, ESI+). Statistical comparisons used Welch’s ANOVA followed by Games–Howell multiple-comparisons testing; *P < 0.05, **P < 0.01; ns, not significant. **e,** Fold change relative to untreated controls for ten of the thirteen targeted measurements, shown with 95% confidence intervals on a log₂ axis; the dashed line denotes no change. Purple, SAIRGA; amber, SAIRGQ. For metabolites measured in both ionization modes, the measurement with narrower confidence intervals is shown; bacillaene, measured with equal precision in both modes, is retained in both, and the three negative-mode surfactin measurements are omitted. Intervals are Welch estimates and are not corrected for multiple comparisons; the Games–Howell values in Supplementary Table 7 are the inferential statistics for all thirteen measurements.

### SAIRGA induces programme-level proteomic remodelling centered on late sporulation

To determine whether the developmental and metabolic phenotypes induced by SAIRGA were accompanied by coordinated changes at the protein level, we performed data-independent acquisition (DIA) proteomics on matrix-associated/extracellular-enriched (F1) and intracellular/cell-associated-enriched (F2) fractions isolated from the same 72-h colony biofilms (Fig. 6; Extended Data Fig. 9). While no single protein met the threshold for statistical significance after Benjamini–Hochberg correction in either fraction, we reasoned that subtle, coordinated shifts across functional groups might still be present. We therefore applied CAMERA-PR across curated biological programmes to identify pathway-level responses.

**Fig. 6.**
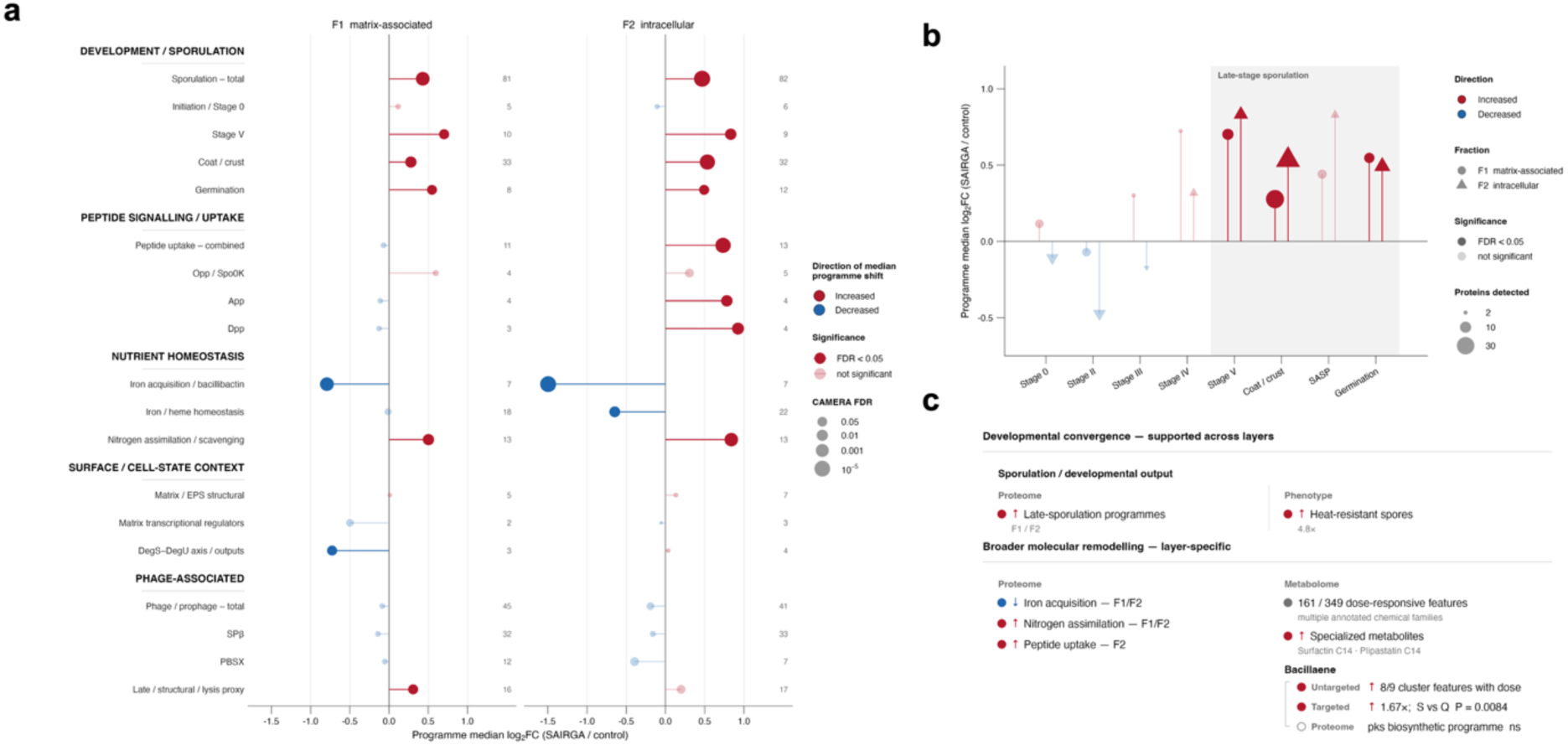
| Program-level proteomic remodeling in SAIRGA-treated *B. subtilis* and convergence with developmental and metabolic phenotypes. Matrix-associated/extracellular-enriched (F1) and intracellular/cell-associated-enriched (F2) fractions from untreated and SAIRGA-treated colony biofilms were analyzed by DIA proteomics (n = 3 biological replicates per condition). Protein intensities were probabilistic-quotient-normalized, log₂-transformed, and analyzed with limma. No individual protein passed Benjamini–Hochberg correction at FDR < 0.05; coordinated responses were therefore assessed by CAMERA-PR across curated functional programs. Program significance is reported as Benjamini– Hochberg-adjusted CAMERA-PR FDR and direction as program median log₂ fold change (SAIRGA versus untreated). **a,** Program landscape in F1 and F2. Point position indicates program median log₂ fold change, color indicates direction, point size reflects −log₁₀(FDR), and saturated symbols denote FDR < 0.05. Numbers indicate the number of detected proteins contributing to each program; indented entries denote nested subprogrammes. **b,** Sporulation programs resolved by developmental stage. Color indicates direction, symbol shape indicates fraction, point size indicates the number of detected proteins, and saturated symbols denote FDR < 0.05. Stage V, coat/crust and germination programs were significantly enriched in both fractions, whereas initiation and early-to-mid sporulation programs were not. **c,** Integration of proteomic, phenotypic and metabolomic responses. Late-sporulation-associated proteomic enrichment is shown alongside the independently measured increase in heat-resistant spores; broader program-level changes are shown together with the untargeted and targeted metabolomic responses. Bacillaene is shown as an example in which metabolite abundance increased without significant coordinated enrichment of the corresponding bacillaene/*pks* proteomic program. Filled red and blue symbols denote increased and decreased responses, respectively; grey denotes a non-directional metabolomic summary, and the open grey symbol denotes no significant program-level change. Measurements derive from separate experiments and analytical layers and are not represented on a common quantitative scale.

The dominant programme-level response was associated with sporulation (Fig. 6a,b). Enrichment was concentrated in late developmental modules rather than at sporulation initiation. Stage V proteins were significantly enriched in both F1 and F2 (FDR = 0.032 and 0.0082, respectively), as were coat/crust proteins (FDR = 9.2 × 10⁻³ and 2.3 × 10⁻⁵) and germination-associated proteins (FDR = 0.037 and 0.031). By contrast, initiation and early-to-mid sporulation programmes showed no significant coordinated enrichment. Protein-level inspection confirmed that these programme-level signals were distributed across multiple late-sporulation and mature-spore-associated proteins rather than being driven by a single outlier (Extended Data Fig. 10b). Binary detection provided additional support for this interpretation, as the germination-associated proteins GerKA and CwlD were detected in all three SAIRGA-treated F1 samples but in none of the untreated samples (Extended Data Fig. 10c). We interpret these detection gains as supportive rather than independent evidence, because treated F1 samples showed a modest overall increase in protein identifications. Together with the programme-level analysis, however, they reinforce the predominance of a mature-spore-associated proteomic state (Extended Data Fig. 10e).

The late-sporulation signature closely matched the phenotype measured independently in intact colony biofilms. SAIRGA increased heat-resistant spore recovery by approximately 4.8-fold without increasing total viable CFU, providing phenotypic support for the increased representation of mature-spore-associated proteins detected by proteomics (Fig. 6c). Because proteomic samples were collected after 72 h of colony development, these data define the endpoint developmental state but do not identify the upstream event that initiated the sporulation decision.

Beyond sporulation, SAIRGA induced broader coordinated remodeling of surface-associated regulatory and metabolic programs. DegS–DegU-associated proteins showed coordinated remodeling, providing a regulatory connection to the surface-associated developmental response. Iron-acquisition/bacillibactin proteins shifted coordinately downward, whereas nitrogen-assimilation/scavenging and peptide-uptake programs shifted upward, with peptide-uptake enrichment particularly evident in F2 (Fig. 6a and Extended Data Fig. 10d). In contrast, total prophage, SPβ and PBSX programs showed no coordinated enrichment, providing no evidence for canonical prophage induction as a dominant component of the proteomic response.

We next integrated the proteomic response with the independently measured developmental and metabolomic phenotypes (Fig. 6c). The clearest convergence was observed for sporulation: enrichment of late-sporulation, coat/crust and germination programs in both proteomic fractions paralleled an approximately 4.8-fold increase in heat-resistant spores, without an increase in total viable cell yield. The metabolome showed a similarly broad response, with 161 of 349 features exhibiting significant dose-dependent trends, while targeted analysis in an independent experiment identified increases in specialized metabolites including surfactin C14 and plipastatin C14.

Changes in bacillaene levels illustrate that correspondence between the proteomic and metabolomic layers was not necessarily one-to-one. Eight of nine features within the bacillaene- and-derivatives molecular-network component increased with SAIRGA dose, and targeted quantification in an independent experiment showed a 1.67-fold increase relative to untreated biofilms (Games–Howell-adjusted P = 0.0042), with a significant difference from SAIRGQ (P = 0.0084). Nevertheless, the corresponding bacillaene/pks proteomic program was not significantly enriched in either F1 or F2 (CAMERA-PR FDR = 0.24 and 0.40, respectively). Thus, the metabolite-level response was not accompanied by detectable coordinated enrichment of the corresponding biosynthetic protein program. The correspondence between molecular layers was therefore strongest at the level of biological state and functional programs rather than through one-to-one coupling between metabolite abundance and biosynthetic protein abundance. Collectively, the proteomic data place the metabolomic changes within a broader developmental response. SAIRGA-treated colony biofilms were characterized most prominently by increased representation of late-sporulation and mature-spore-associated programs, together with remodeling of nutrient- and metabolic-response pathways. This molecular state closely parallels the increase in heat-resistant spore formation, while the lack of coordinated canonical prophage induction indicates that the response is not accompanied by a broad canonical prophage-induction program.

## Discussion

Our findings reveal that the arbitrium peptide SAIRGA can reshape bacterial development outside the phage infection context in which its signaling function is known. Rather than producing a general growth response, SAIRGA altered the organization of a structured *B. subtilis* colony, with spatial redistribution of matrix-associated activity alongside a marked increase in sporulation. This places a phage-derived communication molecule among the cues that can influence bacterial developmental state and expands the biological context in which arbitrium-associated chemistry can be considered.

The combination of spatial *PtapA* remodeling and increased sporulation is central to this interpretation. *B. subtilis* colony biofilms are developmentally heterogeneous communities in which matrix production and sporulation are organized in space and time. Matrix-producing populations can form a propagating front at the colony periphery, followed by a spatially displaced sporulation front as the community develops ^17^. Against this background, the *PtapA* response to SAIRGA is poorly described as simple activation or repression of matrix production. At the colony edge, mean reporter activity increased while high local maxima decreased, producing a broader and less sharply concentrated pattern of activity. At the same developmental endpoint, heat-resistant spore recovery increased approximately 4.8-fold and sporulation efficiency rose from approximately 5% to 19%. Total viable cell yield, however, remained similar between treated and untreated colonies. The larger colony footprint therefore does not reflect a corresponding increase in viable biomass; instead, the spatial and sporulation readouts together indicate a change in how developmental states are represented within the colony.

The strong dependence of this response on growth context may provide a clue to how it is organized. SAIRGA produced a prominent phenotype in surface-grown colonies, whereas planktonic growth remained largely unchanged and static-liquid pellicles did not display a comparable macroscopic response. Colony biofilms and pellicles share major matrix components and regulatory systems but develop under different spatial, mechanical and metabolic conditions ^6,7^. Such differences are relevant to developmental signaling in *B. subtilis*. Mechanical information associated with flagellar restraint can feed into the DegS–DegU system^19^, and DegU activity contributes to transitions among motile, matrix-producing and other biofilm-associated states^20^. In particular, DegU phosphorylation can alter cell-fate distribution in B. subtilis biofilms, reducing the fraction of cells entering matrix-producing states while favoring Spo0A-associated progression towards sporulation^21^.

This makes the DegS–DegU/Spo0A network an intriguing regulatory context for the SAIRGA phenotype. Consistent with this possibility, proteomic analysis identified coordinated remodeling of a DegS–DegU-associated program. The significance of this observation lies less in the abundance of DegU itself—which did not change significantly—than in the biological position of this regulatory axis at the interface between environmental information, matrix-associated states and sporulation. Our data therefore suggest a point of convergence between the phenotype and a known developmental regulatory landscape, rather than defining a linear SAIRGA→DegU→Spo0A pathway. Direct measurements of DegU phosphorylation and genetic perturbation of this network will be needed to determine whether it participates causally in the response.

The sequence and stereochemical requirements of the phenotype further indicate that SAIRGA is not acting simply as an added nutrient or through a nonspecific extracellular effect. The tested N-terminal Ser-to-Gly substitution was tolerated, whereas the C-terminal Ala-to-Gln substitution abolished the colony response, and the all-D enantiomer failed to reproduce the activity of L- SAIRGA. These observations are consistent with a stereoselective step in peptide transport, processing or molecular recognition. Yet the response persisted in a ΔaimR background, demonstrating that the receptor responsible for interpreting SAIRGA in the canonical φ3T arbitrium circuit is dispensable for this bacterial phenotype^11,13,14^. AimR independence therefore separates the developmental response from the canonical arbitrium recognition pathway, while leaving the host component responsible for sensing or processing SAIRGA unresolved.

The ability of *B. subtilis* to respond to SAIRGA also raises a broader question about how chemical information is interpreted across phage–host boundaries. SAIRGA evolved as a signal within phage communication^11^, but host responsiveness does not by itself imply that the peptide evolved to manipulate bacterial development. From the bacterial perspective, the molecule may instead function as a cue: information generated in one biological interaction that can nevertheless be detected and incorporated into another organism’s physiology. Such recognition could arise through overlap with existing peptide-sensing or peptide-transport architectures, or it could represent a more specific capacity to respond to molecules associated with phage activity. Distinguishing these possibilities will require identification of the host machinery responsible for the AimR-independent phenotype.

Recent mechanistic studies of φ3T provide a useful contrast to this host response. During infection, the arbitrium-controlled phage regulatory network recruits the host-encoded MazEF toxin– antitoxin system as part of the machinery governing the phage lysis–lysogeny decision^22,23^. In those studies, bacterial MazEF is used within an infection-associated circuit controlling phage fate; this is distinct from a bacterial developmental response to the mature communication molecule itself. Here, SAIRGA was supplied in the absence of experimentally imposed infection and was sufficient to alter bacterial development without a requirement for AimR. The two settings therefore highlight different ways in which phage communication chemistry can intersect with the host: through host machinery incorporated into a viral life-cycle decision during infection, and through a physiological response of the bacterial community to the mature peptide outside that infection programme.

A broader ecological context comes from recent work showing that phage infection itself can alter bacterial developmental timing. Măgălie and colleagues observed localized early sporulation around advancing infection fronts in spatially structured *B. subtilis* populations, with consequences for both plaque expansion and persistence of phage genomes within spores^24^. Filtered phage lysate was insufficient to reproduce this response, leaving the chemical or cellular trigger unresolved. Notably, our findings point to a complementary phenomenon: in a distinct experimental setting, a defined phage communication peptide supplied independently of infection is sufficient to increase sporulation in a structured bacterial population. Whether arbitrium signalling contributes to developmental responses during infection now becomes an experimentally testable question.

The molecular profiling further shows that this developmental response is embedded within a broader physiological transition. Untargeted metabolomics revealed a graded response across the SAIRGA concentration series, and the relationship to peptide activity was reproduced across sequence variants: the active peptide GAIRGA generated a strongly concordant metabolic response, whereas the inactive C-terminal variant SAIRGQ produced little comparable perturbation. The metabolome therefore provides an independent indication that the colony phenotype reflects a change in physiological state rather than morphology alone.

Within this response, bacillaene provides the clearest specialized-metabolite anchor. Most features within the bacillaene molecular family increased with SAIRGA dose, and targeted measurements in an independently grown experiment reproduced the increase and distinguished SAIRGA from SAIRGQ. Selected surfactin and plipastatin congeners showed more modest changes in the same direction. Specialized-metabolite production is closely integrated with differentiation in *B. subtilis* communities^10^, making these changes compatible with a shift in differentiated colony physiology. Their role should not, however, be reduced to direct pathway activation: the bacillaene metabolite response was not accompanied by significant coordinated enrichment of the corresponding *pks* protein programme. Instead, bacillaene appears here as a reproducible chemical output of the altered state.

Other metabolomic changes suggest that the transition extends beyond specialized metabolism. Tryptophan-related chemistry displayed a structured bidirectional response, with a tryptophan/indole-associated molecular family decreasing while structurally distinct tryptophan-derived amide- and peptide-like features increased. Short-peptide and lipid-associated families also showed directional changes. At the protein level, nitrogen assimilation/scavenging was coordinately remodelled, and peptide-uptake programmes were strongly increased in the cell-associated fraction, while iron-acquisition machinery shifted in the opposite direction. Taken together, these observations suggest altered nutrient and peptide handling alongside developmental reorganization, although the present endpoint measurements do not establish whether these metabolic changes precede the developmental shift or arise as part of the differentiated state that follows it. Notably, this peptide-uptake signature coincided with very low recovery of exogenously supplied SAIRGA from 72-h colony-containing samples.

Proteomics most clearly converged with the direct phenotype at the level of sporulation. The dominant coordinated response was concentrated in Stage V, coat/crust and germination-associated programmes, closely matching the increased recovery of heat-resistant spores from intact colonies. The proteomic profile therefore captures the altered developmental composition reached by SAIRGA-treated communities at the 72-h endpoint and provides an independent molecular counterpart to the sporulation phenotype. More broadly, the partial separation between proteomic and metabolomic responses—including the bacillaene example—supports interpretation of the omics layers as complementary views of a changed physiological state rather than components of a single linear signalling pathway.

Several important questions now follow directly from this framework. The host machinery responsible for the AimR-independent response remains unknown, including whether stereoselectivity arises during peptide uptake or molecular recognition. The temporal relationship between spatial *PtapA* reorganization and commitment to sporulation also remains to be resolved. Identifying the relevant peptide-handling or recognition machinery, following developmental reporters through time, and perturbing candidate DegS–DegU/Spo0A-associated pathways should distinguish events involved in sensing SAIRGA from downstream consequences of the altered colony state. A further intriguing question is whether the developmental response described here influences phage infection itself. Introducing defined arbitrium peptide perturbations into spatial infection models could test whether the host response feeds back on plaque propagation, lysogeny or persistence within spores.

Taken ogether, our results reveal a bacterial developmental response to a molecule whose established signalling function lies in communication between phages. In a structured B. subtilis colony, SAIRGA is sufficient, outside experimentally imposed infection, to alter matrix-associated spatial fluorescence, increase sporulation and shift the broader physiological state of the community. The colony phenotype persists in the absence of AimR, placing this macroscopic response outside the canonical AimR-dependent recognition pathway. These findings extend the biological context of arbitrium-associated chemistry beyond the viral life-cycle decision and show how molecules produced for phage communication can become part of the chemical information landscape that shapes bacterial development.

## Methods

### Bacterial strains and growth conditions

*Bacillus subtilis* NCIB 3610 was used throughout. Two derivative strains were used: a Δ*aimR* deletion strain and a P*tapA*–*yfp* transcriptional reporter strain. Both strains were obtained from external sources. Full genotypes, strain designations and provenance will be reported on publication.

For each experiment, B. subtilis NCIB 3610 was streaked from a frozen glycerol stock onto an LB agar plate and grown overnight at 37 °C. Starter cultures were prepared by transferring a loopful of growth from the plate into 2 mL Luria–Bertani (LB) medium (10 g L⁻¹ tryptone, 5 g L⁻¹ yeast extract and 10 g L⁻¹ NaCl) and growing for 4 h at 37 °C with shaking at 220 rpm.

Biofilm-inducing MSgg medium was prepared essentially as described previously ^8^ and contained 5 mM potassium phosphate (pH 7.0), 100 mM MOPS (pH 7.0), 2 mM MgCl₂, 700 µM CaCl₂, 50 µM MnCl₂, 50 µM FeCl₃, 1 µM ZnCl₂, 2 µM thiamine, 0.5% (v/v) glycerol, 0.5% (w/v) monosodium glutamate and 50 µg mL⁻¹ each of phenylalanine, tryptophan and threonine. FeCl₃ was added immediately before use. For solid medium, agar was autoclaved separately and combined with 2× MSgg at approximately 55 °C to a final agar concentration of 1.5% (w/v). Plates (60 mm) were poured and dried overnight in the dark. Planktonic growth assays were performed at 37 °C; colony biofilm and pellicle assays were performed at 30 °C.

### Peptide synthesis and characterization

SAIRGA (Ser-Ala-Ile-Arg-Gly-Ala), GAIRGA, SAIRGQ and GMPRGA were synthesized in-house by manual solid-phase peptide synthesis using a standard Fmoc/t-Bu strategy at 0.2 mmol scale on Rink Amide MBHA resin (0.74 mmol g⁻¹), yielding C-terminally amidated peptides. Fmoc deprotection was performed with 20% (v/v) piperidine in DMF (2 × 8 min), and Fmoc-amino acids were coupled in fourfold excess with HBTU/DIPEA (1:1:20) in DMF for 90 min per residue. Peptides were cleaved with TFA/triisopropylsilane/water (95:2.5:2.5, v/v/v) for 2 h at room temperature.

Purification was performed by preparative reverse-phase HPLC (Dionex Ultimate 3000; ZORBAX 300SB-C18 PrepHT, 21.2 × 250 mm, 7 µm; Agilent) at 20 mL min⁻¹ using solvent A (99% water, 1% acetonitrile, 0.1% TFA) and solvent B (90% acetonitrile, 10% water, 0.07% TFA). Purity was assessed by analytical HPLC (ZORBAX 300SB-C18, 4.6 × 150 mm, 5 µm) using the same solvent system. Only preparations exceeding 95% purity were used, and peptide identity was confirmed by ESI-HRMS (Supplementary Figs. 1 and 3).

D-SAIRGA, the all-D enantiomer of SAIRGA, was obtained commercially (GL Biochem (Shanghai) Ltd, Shanghai, China). All other peptides were L-enantiomers. Peptides were prepared as C-terminal amides and are referred to by sequence throughout; the same C-terminal chemistry was used for all sequence variants and stereoisomers.

GAIRGA and SAIRGQ were designed as sequence controls. GAIRGA carries an N-terminal Ser1→Gly substitution while retaining the C-terminal RGA motif, whereas SAIRGQ carries an Ala6→Gln substitution designed to perturb the C-terminal region implicated in AimR–peptide recognition. GMPRGA is the arbitrium peptide associated with the endogenous SPβ prophage of NCIB 3610. D-SAIRGA was used as a stereochemical probe of peptide uptake or molecular recognition.

Peptides were dissolved in sterile distilled water and added to media at the indicated final concentrations. In an initial planktonic-growth experiment, SAIRGA was prepared in DMSO; matched vehicle controls were included, and DMSO concentrations up to 2% (v/v) did not detectably alter growth.

### Planktonic growth assays

Peptides were added to clear-bottom 96-well microtiter plates containing LB or MSgg at final concentrations of 0, 0.78, 1.56, 3.13, 6.25, 12.5, 25, 50, 100 and 200 µM. Starter cultures were diluted to a starting OD₆₀₀ of 0.05. Plates were incubated at 37 °C in a Varioskan LUX multimode microplate reader (Thermo Fisher Scientific), and OD₆₀₀ was recorded every 20 min. Three independent biological replicates were analyzed per condition. The 0–8 h interval and concentrations up to 100 µM are shown in the figures; the 200 µM condition was retained in the source data. SAIRGA growth assays were performed in two independent experiments. The earlier experiment used DMSO stocks with matched vehicle controls; subsequent experiments used peptide stocks prepared in sterile water.

### Colony biofilm assays and morphometric analysis

MSgg agar and medium components were combined at no more than 55 °C. Peptides were added after mixing, plates were poured immediately following brief vortexing and were dried overnight under sterile conditions in the dark. A 3 µL droplet of freshly prepared starter culture was spotted at the center of each 60-mm plate, and plates were incubated at 30 °C for 72 h before imaging. SAIRGA was tested at 1, 5, 10, 20 and 50 µM; concentrations used in individual experiments are specified in the corresponding figure legends. Three independent biological replicates were analyzed per condition.

The experiments comparing peptide sequence, stereochemistry and AimR dependence were performed as three independent biological experiments, each with its own contemporaneous untreated control: (i) wild-type untreated, SAIRGA, GMPRGA and GAIRGA; (ii) wild-type untreated, SAIRGA and D-SAIRGA; and (iii) Δ*aimR* untreated and SAIRGA. Absolute colony areas were therefore compared within experiments and not across panels.

Colony area was quantified in Fiji/ImageJ from top-view images acquired under comparable conditions. Each image was calibrated to the internal plate diameter (60 mm), and the colony boundary was manually delineated as the outermost continuous region of visible bacterial growth, including low-density peripheral expansion in peptide-treated colonies.

### Pellicle assays

For pellicle formation, 3 µL starter culture was inoculated into 3 mL MSgg in glass tubes in the absence or presence of 10 µM SAIRGA and incubated statically at 30 °C for 72 h. Pellicles were imaged from above and from the side. For crystal-violet staining, unattached cells and medium were removed, biofilms were stained with 0.1% (w/v) crystal violet for 15 min and washed twice with 0.1% (w/v) NaCl. Crystal-violet staining was assessed qualitatively and was not quantified spectrophotometrically.

### Sporulation and viable-cell count

Colony biofilms were grown on MSgg agar at 30 °C for 72 h in the absence or presence of SAIRGA. Each whole colony together with the surrounding biofilm was harvested from the plate into 10 mL PBS containing 0.05–0.1% (v/v) Tween-80 by gentle scraping until no visible biomass remained. Biofilms were dispersed by vortexing (3 × 1 min, with 30 s rests), followed by addition of EDTA to 2 mM final concentration, bath sonication in an ice-water bath (3 × 1 min at standard power, with 30 s rests and brief vortexing between cycles), and a final 30 s vortex with vigorous pipetting; total EDTA contact time was kept below approximately 10 min. A 1:100 working stock was prepared in fresh Tween-PBS without EDTA, and serial ten-fold dilutions were prepared from this stock. For total viable CFU, 100 µL of the appropriate dilution was spread-plated. To determine heat-resistant CFU, a 1 mL aliquot of the same working stock was heat-shocked by placing in a water bath at 80 °C for 10 min, cooled on ice for 5 min, serially diluted and plated identically. Plates containing 30–300 colonies were counted after incubation at 30 °C for 16–24 h.

Three independent biological replicates were analysed per condition; for each biological replicate, counts were averaged from three technical plating replicates. Total and heat-resistant CFU were compared using unpaired t-tests with Welch’s correction on log10-transformed CFU. Sporulation efficiency was calculated for each biological replicate as (heat-resistant CFU / total viable CFU) × 100 and was reported descriptively as a derived ratio.

### Fluorescence imaging and spatial analysis of PtapA activity

Colony biofilms of the P*tapA*-YFP transcriptional reporter strain and the non-fluorescent wild-type control were grown on MSgg agar (1.5% w/v) at 30 °C for 72 h. Where indicated, SAIRGA was incorporated into the medium at 20 µM during plate preparation, 24 h before inoculation. Untreated reporter biofilms and the non-fluorescent wild-type strain served as controls. Three independent colony biofilms were analyzed per condition.

Biofilm morphology and fluorescence were imaged using a Leica M165 FC stereomicroscope. All images used for quantitative comparison were acquired with identical gain settings (2.19); exposure times were 4.44 s for whole-biofilm measurements and 8.6 s for zonal measurements. Fluorescence was quantified in Fiji/ImageJ. Three radial zones were defined: edge, intermediate and center. For zonal analysis, a fixed rectangular region of interest (181,760 pixels) was used to record mean and maximum fluorescence intensity. Edge and intermediate zones were sampled in two spatial fields per biofilm, and the center once per biofilm, with two technical ROI measurements per field. Spatial fields and technical ROI measurements were treated as nested subsamples; the colony biofilm was the biological replicate.

For radial-intensity analysis, two line profiles were extracted per biological replicate from the colony periphery toward the center. Profiles were aligned to the edge-associated fluorescence peak. Because profile lengths ranged from 777 to 963 pixels, profiles were truncated at 700 pixels for visualization and averaging so that all six profiles per condition contributed across the plotted range. Radial profiles were displayed descriptively as mean ± s.d.; the biological replicate remained the colony biofilm (n = 3 per condition).

### Metabolomics experimental design

Two independent metabolomics experiments were performed. Experiment 1 was an untargeted dose-response experiment in which *B. subtilis* NCIB 3610 colony biofilms were grown on MSgg agar for 72 h at 30 °C with SAIRGA or GAIRGA at 1, 5, 10 or 50 µM, alongside untreated controls. Five independent biological replicates were analyzed per condition (n = 5). This experiment provided the primary SAIRGA dose-response dataset and the SAIRGA–GAIRGA comparison. Peptide-containing agar plates without bacterial inoculation were prepared at 1, 5, 10 and 50 µM and extracted identically, providing a reference for peptide recovery from the medium.

Experiment 2 was an independent specificity and targeted follow-up experiment comparing 20 µM SAIRGA, 20 µM SAIRGQ and untreated controls. Six independent biological replicates were analyzed per group (n = 6). The same samples were used for targeted quantification and for an untargeted negative-ionization analysis of peptide specificity.

### Metabolite extraction

For both experiments, the entire agar plate, including the colony biofilm and surrounding agar, was transferred to a 50 mL tube. HPLC-grade methanol was added until the agar was fully submerged. Samples were vortexed vigorously, incubated at room temperature for 30 min, vortexed again and centrifuged at 4,000 × g for 20 min at 4 °C. The supernatant was collected, mixed 1:1 (v/v) with LC-MS-grade water and filtered through a 0.22 µm filter directly into an HRMS vial. Samples were stored at −20 °C until analysis; injection volume was 20 µL.

#### Experiment 1: untargeted LC–HRMS/MS acquisition

Untargeted metabolomic profiling was performed on a Q Exactive Focus hybrid quadrupole-Orbitrap mass spectrometer (Thermo Fisher Scientific) coupled to a Dionex Ultimate 3000 UHPLC system. Chromatographic separation used an Accucore C18 reversed-phase column (2.6 µm, 100 × 2.1 mm; Thermo Fisher Scientific) at 40 °C. Mobile phase A was water with 0.1% (v/v) formic acid and mobile phase B was acetonitrile with 0.1% (v/v) formic acid. The gradient was 0– 2 min, 100% A; 2–4 min, linear ramp to 50% B; 4–11 min, linear ramp to 90% B; 11–12 min, 90% B; 12–13 min, linear decrease to 20% B; 13–14 min, 20% B, at 0.4 mL min⁻¹.

Data were acquired in positive ionization mode using a heated electrospray ionization source (spray voltage 3,800 V; capillary temperature 300 °C; sheath gas 35 arbitrary units; S-lens RF level 50). Full MS1 spectra were acquired at 70,000 resolution over *m/z* 150–1,700 (AGC target 1 × 10⁶). Data-dependent MS2 fragmentation was performed on the three most abundant precursor ions per scan at 35,000 resolution, with a 3.0 *m/z* isolation window and stepped normalized collision energies of 25, 30 and 35 eV. Full-MS intensities provided quantitative feature data; discovery MS2 spectra were used for molecular networking and metabolite annotation.

#### Experiment 2: specificity and targeted LC–HRMS acquisition

Experiment 2 used the same instrument and column with an extended 21-min chromatographic program optimized for large lipopeptides: 0–2 min, 100% A; 2–14 min, linear ramp to 95% B; 14– 17 min, 95% B; 17–17.2 min, linear decrease to 0% B; 17.2–21 min, re-equilibration at 100% A; flow rate 0.4 mL min⁻¹. Samples were acquired in both positive and negative ionization modes. Positive-mode source parameters were: spray voltage 2,800 V; capillary temperature 320 °C; sheath gas 50 arbitrary units; auxiliary gas 15 arbitrary units; probe heater temperature 400 °C; S-lens RF level 50. In negative mode, spray voltage was 3,200 V, capillary temperature 320 °C and S-lens RF level 50. Full MS1 spectra were acquired at 70,000 resolution over *m/z* 100–1,500 (AGC target 1 × 10⁶).

Biological samples were acquired in Full MS mode for quantitative analysis. Negative-mode Full-MS data were also processed as an untargeted dataset for comparison of SAIRGA, SAIRGQ and untreated samples. A pooled quality-control sample was injected every five biological injections.

#### Internal standards and negative-mode reinjection

Internal standards were added at 10 µM before injection. Positive-mode targeted measurements of bacillaene and surfactin homologs were normalized to terfenadine ([M+H]⁺, *m/z* 472.321; coefficient of variation 0.34–0.60% across samples; one-way ANOVA across treatment groups, *P* = 0.53). For the final negative-mode analysis, the same Experiment 2 extracts were re-injected after addition of chlorpropamide ([M−H]⁻, *m/z* 275.026; retention time approximately 8.8 min). Chlorpropamide was stable across the negative-mode run (overall CV 10.7%; one-way ANOVA across treatment groups, *P* = 0.16) and was used to normalize all final negative-mode targeted measurements. An earlier candidate internal standard, erythromycin, was rejected because of unstable recovery (CV 31% in control samples). Only the chlorpropamide-normalized reinjection was carried forward for the final negative-mode targeted analysis.

### Additional MS/MS acquisition for annotation

Following the initial untargeted analysis, two selected stored Experiment 1 extracts were reanalyzed using inclusion lists designed to acquire additional MS2 spectra for prioritized features lacking informative fragmentation. These acquisitions were used solely to increase spectral coverage for annotation and did not contribute quantitative abundance measurements.

### Untargeted metabolomics data processing

Raw files were converted to centroided mzML format using MSConvert (ProteoWizard). Feature detection and alignment for the primary Experiment 1 dataset were performed in MZmine 3 (v3.1.0). Chromatograms were built with the ADAP chromatogram builder (*m/z* tolerance 0.002 Da or 10 ppm) and resolved using the local minimum resolver (retention-time search range 0.05 min; peak duration 0–1.0 min). Isotope grouping used a tolerance of 0.001 Da or 5 ppm with a retention-time tolerance of 0.02 min. Features were aligned with the join aligner (*m/z* tolerance 0.001 Da or 5 ppm; retention-time tolerance 0.06 min; *m/z* weight 3.0; retention-time weight 1.0) and gap-filled with matching tolerances.

Feature detection used a minimum peak-height threshold of 5 × 10⁵ and yielded 606 features. Filtering against procedural blanks using a sample-to-blank ratio >1.3 retained 434 features. Suspected chromatographic peak-splitting artifacts were audited by independent reintegration of features sharing accurate mass (±0.005 Da) in contiguous retention-time ladders using revised peak-resolution parameters. Series that collapsed on reintegration were classified as splitting artifacts, whereas reproduced series were retained as genuine isomers. This audit removed 82 splitting artifacts, two features with no detections across the 25-sample dataset and one additional chromatographic artifact confirmed by targeted reinjection, yielding 349 high-confidence features for quantitative analysis. Reintegration was used to adjudicate suspected artifacts; quantitative values for retained features were taken from the primary processing pipeline. Per-feature filtering decisions are provided in the source data.

Peptide-derived features were identified by accurate mass and early retention time (0.47 min) and were excluded from the quantitative feature table.

Non-detected values were retained as NA and, after log₂ transformation, were imputed separately within each sample by random draws from a down-shifted normal distribution. For each sample, the imputation distribution was centered at the mean observed log₂ intensity minus 1.6 times its standard deviation, with a standard deviation of max(0.3 × s.d., 0.1). A fixed random seed (42) was used. Probabilistic quotient normalization (PQN) was then applied to the completed log₂ matrix.

### Statistical analysis of untargeted metabolomics

Global separation among treatment groups in the primary SAIRGA dose-response dataset was assessed by PERMANOVA (adonis2, vegan package) on Euclidean distances of the normalized log₂ matrix using 999 permutations, followed by pairwise comparisons of each SAIRGA concentration with untreated controls. Principal component analysis was performed on the same matrix without scaling. Differential abundance was assessed by Welch’s *t*-test comparing each concentration with untreated controls, followed by Benjamini–Hochberg correction (FDR < 0.05). Dose-response trends were tested by Spearman rank correlation with Benjamini–Hochberg correction.

As a sensitivity analysis during chromatographic curation, non-detected values were also represented using a censoring-aware rank encoding in which non-detects were tied below all observed intensities without assigning numerical abundances. Applied to the immediately preceding 350-feature curation state, this analysis recovered 157 of the 161 trends retained in the final analysis, with agreement between rank-correlation coefficients of r = 0.987.

For the direct SAIRGA–GAIRGA comparison, samples from Experiment 1 were processed in a shared feature table using identical parameters and the same untreated controls. This shared table was used only for the between-peptide comparison and was distinct from the final 349-feature curated SAIRGA dose-response table.

For the Experiment 2 specificity dataset, procedural blanks were used for background filtering and were then excluded, together with QC injections, from the biological analysis matrix. Missing values were handled using the same down-shifted-normal imputation framework, and PQN was calculated from the 18 biological samples only. Per-feature group comparisons used Welch’s *t*-tests with Benjamini–Hochberg correction. Global enrichment of nominally significant features (*P* < 0.05) was assessed using 5,000 label permutations. Kolmogorov–Smirnov statistics and Q–Q plots were retained as descriptive assessments of *P*-value distributions, whereas the permutation test provided the primary global inference.

### Molecular networking and metabolite annotation

Feature-based molecular networking was performed on the GNPS2 platform using MS2 spectra from Experiment 1. Networks were generated with a cosine-similarity threshold of 0.7, precursor-ion mass tolerance of 0.02 Da and a minimum of six matched fragment ions per edge, yielding 605 nodes, of which 471 were connected to at least one neighbor. Spectral-library searching against GNPS2 libraries provided library-based annotations.

Molecular formulas, candidate structures and chemical classes were additionally predicted with SIRIUS using ZODIAC, CSI:FingerID and CANOPUS/NPClassifier. GNPS library evidence, SIRIUS structural and formula predictions, CANOPUS class assignments and additional inclusion-list MS2 spectra were integrated at the feature/network-node level. Annotation confidence was assigned as MSI level 2 for GNPS spectral-library matches, MSI level 3 for SIRIUS-only candidate structures or formula assignments supported by a predicted chemical class, and MSI level 4 for molecular-formula assignments without additional class or spectral evidence. Likely contaminants, including buffer-, detergent-, polymer- and synthetic-reagent-related features, were excluded from biological interpretation. Networks were visualized in Cytoscape.

### Targeted metabolomics data processing and statistics

Targeted metabolites were quantified by extracted-ion chromatogram peak-area integration in Xcalibur FreeStyle (Thermo Fisher Scientific). Normalized peak areas were log₁₀-transformed before statistical analysis. Bacillaene and surfactin C13–C15 congeners were quantified in both ionization modes. Where a compound was measured in both modes, the measurement carried forward for the graphical summary was selected according to within-group analytical precision pooled across treatment groups; bacillaene showed comparable precision in both modes and was retained as an orthogonal measurement in each.

Group comparisons were performed in GraphPad Prism v11.0.2 using Welch’s ANOVA followed by Games–Howell multiple-comparisons testing for SAIRGA versus untreated, SAIRGQ versus untreated and SAIRGA versus SAIRGQ. Fold changes were expressed on the linear scale as the back-transformed difference of log₁₀ group means. Six independent biological replicates were analyzed per treatment group (n = 6).

### Proteomic sample preparation and fractionation

For proteomic profiling, colony biofilms were grown in the absence or presence of 20 µM SAIRGA, with three independent biological replicates per condition. Biofilm biomass was collected with inoculating loops without disturbing the agar and resuspended in 200 µL extraction buffer containing 2 M NaCl, 0.5% (w/v) SDS and 1× protease inhibitor cocktail (Halt Protease Inhibitor Cocktail, Thermo Fisher Scientific). Samples were disrupted by bath sonication for 5 min followed by shaking for 5 min.

Samples were centrifuged at 3,000 × g for 10 min at 4 °C. The supernatant was collected as the matrix-associated/extracellular-enriched fraction (F1). The pellet was processed to recover the intracellular/cell-associated-enriched fraction (F2) by probe sonication on ice (seven cycles of 5 s on/10 s off at 20% amplitude), followed by centrifugation at 12,000 × g for 15 min at 4 °C. Protein concentration in each fraction was measured by BCA assay (Pierce Dilution-Free Rapid Gold BCA Protein Assay, Thermo Fisher Scientific), and samples were adjusted to 200 µL at 2 mg mL⁻¹ protein. F1 and F2 were fractions of the same biological samples and were not treated as independent biological replicates.

### SP3-based proteomic clean-up and digestion

Hydrophilic and hydrophobic carboxylate-functionalized paramagnetic beads (Sera-Mag SpeedBeads, Cytiva; 40 µL of each bead type per sample) were combined 1:1 and washed three times with molecular-grade water. Beads were added to each sample and incubated with shaking at 1,000 rpm for 10 min at room temperature. Protein binding was promoted by addition of ethanol to >55% (v/v), followed by a further 10 min at 1,000 rpm; beads were then washed three times with 80% ethanol on a magnetic stand.

Bound proteins were reduced with DTT (200 mM stock, 10 µL) in 0.5% SDS/2 M urea/PBS for 15 min at 65 °C and alkylated with iodoacetamide (400 mM stock, 10 µL) for 30 min at 37 °C in the dark. Proteins were rebound to the beads with ethanol, washed three times with 80% ethanol and resuspended in 150 µL 2 M urea/PBS. Trypsin (Promega, mass-spectrometry grade; 2 µg) and CaCl₂ (4 mM final concentration) were added, and samples were digested overnight at 37 °C with shaking. Peptides were washed with acetonitrile, eluted from the beads in two sequential 100 µL aliquots of 2% DMSO (30 min and 45 min at 37 °C, 1,000 rpm), desalted using C18 tips (Pierce, Thermo Fisher Scientific), dried under vacuum and resuspended in 10 µL 0.1% formic acid. Sample preparation followed established SP3 protocols^25,26^.

### Proteomic LC–MS/MS analysis and DIA-NN processing

All solvents were LC-MS grade (BioLab, Israel). Peptide samples were diluted tenfold with 0.1% TFA, and 3 µL was injected. Samples were loaded using a nano-UPLC system (nanoElute 2, Bruker, Germany) with mobile phase A (water with 0.1% formic acid) and mobile phase B (acetonitrile with 0.1% formic acid). Peptides were desalted on a trap column (PepMap, 100 µm × 5 cm; Thermo Fisher Scientific) and separated on an Aurora analytical column (75 µm i.d. × 25 cm; IonOpticks) at 0.3 µL min⁻¹, using the following gradient: 2% to 30% B over 30 min, 30% to 95% B over 0.5 min and 95% B held for 6.5 min. The nano-UPLC was coupled online to a quadrupole time-of-flight mass spectrometer (timsTOF Ultra, Bruker) through a CaptiveSpray2 nanoESI source (Bruker Daltonics, Germany).

Data were acquired in parallel accumulation–serial fragmentation combined with data-independent acquisition (DIA-PASEF) mode. The MS1 mass range was 100–1,700 m/z and the MS1 ion-mobility range was 0.70–1.30 V s cm⁻² (1/K₀), with a ramp time of 166 ms and an estimated cycle time of 1.2 s. Twenty-seven isolation windows of 25 Da with 1 Da overlap were set over the range 374–1,050 m/z^27^.

DIA data were processed with DIA-NN v2.2.0. Peptide length was restricted to 7–30 amino acids, peptide mass to 100–1,700 Da and precursor charge to 1–4. Up to two missed cleavages were allowed. Carbamidomethylation of cysteine (+57.0215 Da) was specified as a fixed modification; methionine oxidation (+15.9949 Da) and peptide N-terminal acetylation (+42.0106 Da) were variable modifications. FDR was controlled at 1%.

Proteomic sample preparation was carried out at the Proteomics Unit, Ilse Katz Institute for Nanoscale Science and Technology, Ben-Gurion University of the Negev. Prepared samples were transferred to the de Botton Institute for Protein Profiling (G-INCPM, Weizmann Institute of Science), where LC–MS/MS acquisition and DIA-NN protein-group assignment were performed as a service. All downstream processing and statistical analysis reported here — normalization, differential-abundance modelling, program-level enrichment and detection-pattern analysis — were performed by the authors.

### Proteomics data processing and statistical analysis

Protein-group abundance matrices were analyzed separately for F1 and F2. Non-positive intensity values were treated as not detected. PQN was performed independently within each fraction. For each protein, a reference abundance was calculated as the median intensity across the six samples, and sample-specific normalization factors were calculated as the median quotient between sample intensities and corresponding reference values. Normalized intensities were log₂-transformed. Missing values were retained as NA and were not imputed.

Proteins entered quantitative analysis if detected in at least two of three biological replicates in both untreated and SAIRGA-treated groups. Remaining missing observations were retained as NA, and limma linear models were fitted using available non-missing measurements. Differential abundance was modeled separately for F1 and F2 in R (R v4.5.2, Bioconductor v3.22, limma v3.66.0) for the contrast SAIRGA-treated versus untreated. Moderated statistics were obtained using empirical-Bayes moderation with eBayes(trend = TRUE, robust = TRUE), and protein-level *P* values were adjusted using the Benjamini–Hochberg procedure.

### Program-level proteomic analysis

Proteins were assigned to functional programs programmatically using a fixed rule set applied to gene names and protein descriptions. Assignment rules combined standard *B. subtilis* gene-name prefixes (for example *spo*, *cot*, *ssp*, *ger*, *opp*, *app*, *dpp*, *rap*, *phr*, *dhb*, *gln*, *srf*, *pks*, *pps* and *xkd*) with keyword matching in protein descriptions. Programs included sporulation and sporulation- stage modules, spore coat and germination proteins, oligopeptide transport systems (Opp/Spo0K, App and Dpp), Rap and Phr regulators, iron acquisition and bacillibactin-associated proteins, nitrogen assimilation and scavenging proteins, matrix- and motility-associated proteins, secondary-metabolite biosynthetic programs and SPβ- and PBSX-associated prophage proteins. Proteins could be assigned to more than one program. The rule set was defined independently of the differential-abundance results and applied identically to F1 and F2. Complete program membership is provided in the source data (Source Data Fig. 6, sheet S20D_CAMERA_membership).

Program-level enrichment was assessed separately for F1 and F2 using cameraPR from the limma package, with moderated *t*-statistics from the differential-abundance analysis as the ranked input statistic (use.ranks = FALSE, inter.gene.cor = 0.01). Positive statistics indicated higher abundance in SAIRGA-treated samples and negative statistics higher abundance in untreated samples. Program-level *P* values were adjusted by the Benjamini–Hochberg procedure, with FDR < 0.05 considered significant.

### Protein-detection analysis

A descriptive presence/absence analysis was performed in parallel on the unnormalized DIA abundance matrices, without normalization or imputation. For each protein, the number of biological replicates in which it was detected was determined separately for untreated and SAIRGA-treated samples. Detection gains were defined as changes from 0/3 to 2/3, 0/3 to 3/3 or 1/3 to 3/3 detections, with corresponding reverse patterns defined as detection losses. Because n = 3 biological replicates were available per condition, detection switches were used descriptively rather than as individual inferential tests.

Per-sample protein-detection depth and overall gain/loss counts were examined as background calibration. Quantitative abundance and detection status were treated as two readouts of the same DIA measurements, not as independent evidence layers. No missing values were imputed at any stage of the final proteomics workflow.

### Statistical analysis of phenotypic assays

Colony biofilm area was analyzed in GraphPad Prism v11.0.2 using Welch’s ANOVA followed by Games–Howell multiple-comparisons testing on raw area values. Total and heat-resistant CFU were analyzed after log₁₀ transformation using unpaired *t*-tests with Welch’s correction. Sporulation efficiency was reported descriptively as a derived ratio. Zonal P*tapA*-YFP fluorescence was analyzed using a mixed-effects model followed by Šídák multiple-comparisons testing of untreated versus SAIRGA-treated biofilms within the edge, intermediate and center zones. Three independent biological replicates; nested spatial fields and technical ROI measurements were not treated as independent observations. Whole-biofilm fluorescence measurements were retained as descriptive supporting analysis at the biofilm level.

## Data and code availability

Metabolomics feature tables, normalized matrices, per-feature statistics, curation decisions, network node attributes and the source data underlying all figures are provided with the manuscript. Raw mass-spectrometry data will be deposited in public repositories on publication. Analysis scripts are available from the corresponding author on request.

## Acknowledgements

We are grateful to Prof. Ilana Kolodkin-Gal for providing bacterial strains used in this study and for valuable discussions throughout the course of this work.

---

**Extended Data Fig. 1.**
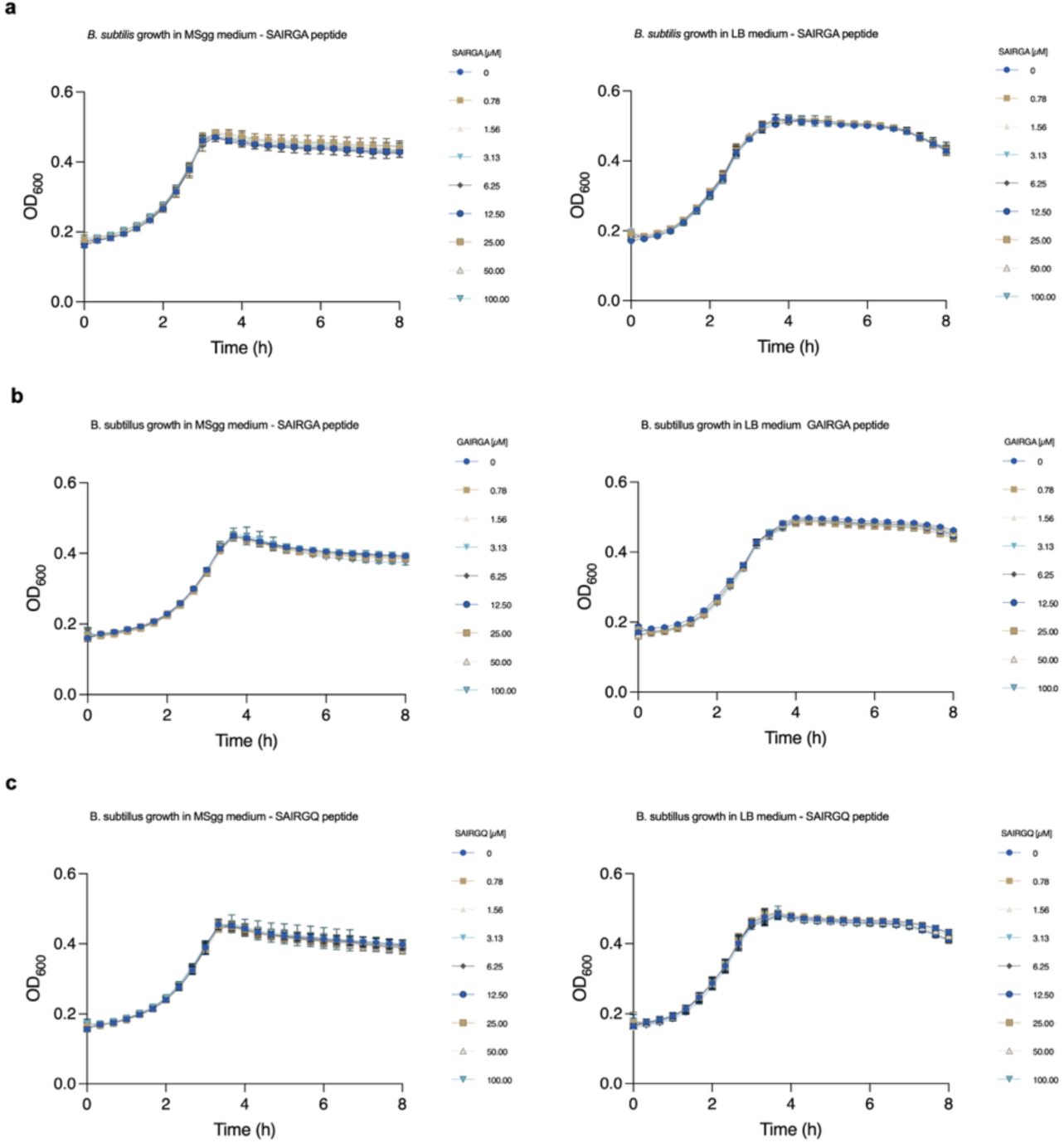
| Planktonic growth kinetics remain similar across SAIRGA and sequence-control peptide concentrations. Growth of *B. subtilis* NCIB 3610 in liquid MSgg or LB supplemented with the indicated concentrations of a, SAIRGA, b, GAIRGA or c, SAIRGQ (MSgg, left; LB, right). OD₆₀₀ was recorded at 20-min intervals at 37 °C in a Thermo Scientific Varioskan LUX microplate reader; the first 8 h, spanning lag phase, exponential growth and stationary phase, are shown. Data are mean ± s.d. of three independent biological replicates. No reproducible concentration-dependent separation of the growth trajectories was observed over the interval shown.

**Extended Data Fig. 2.**
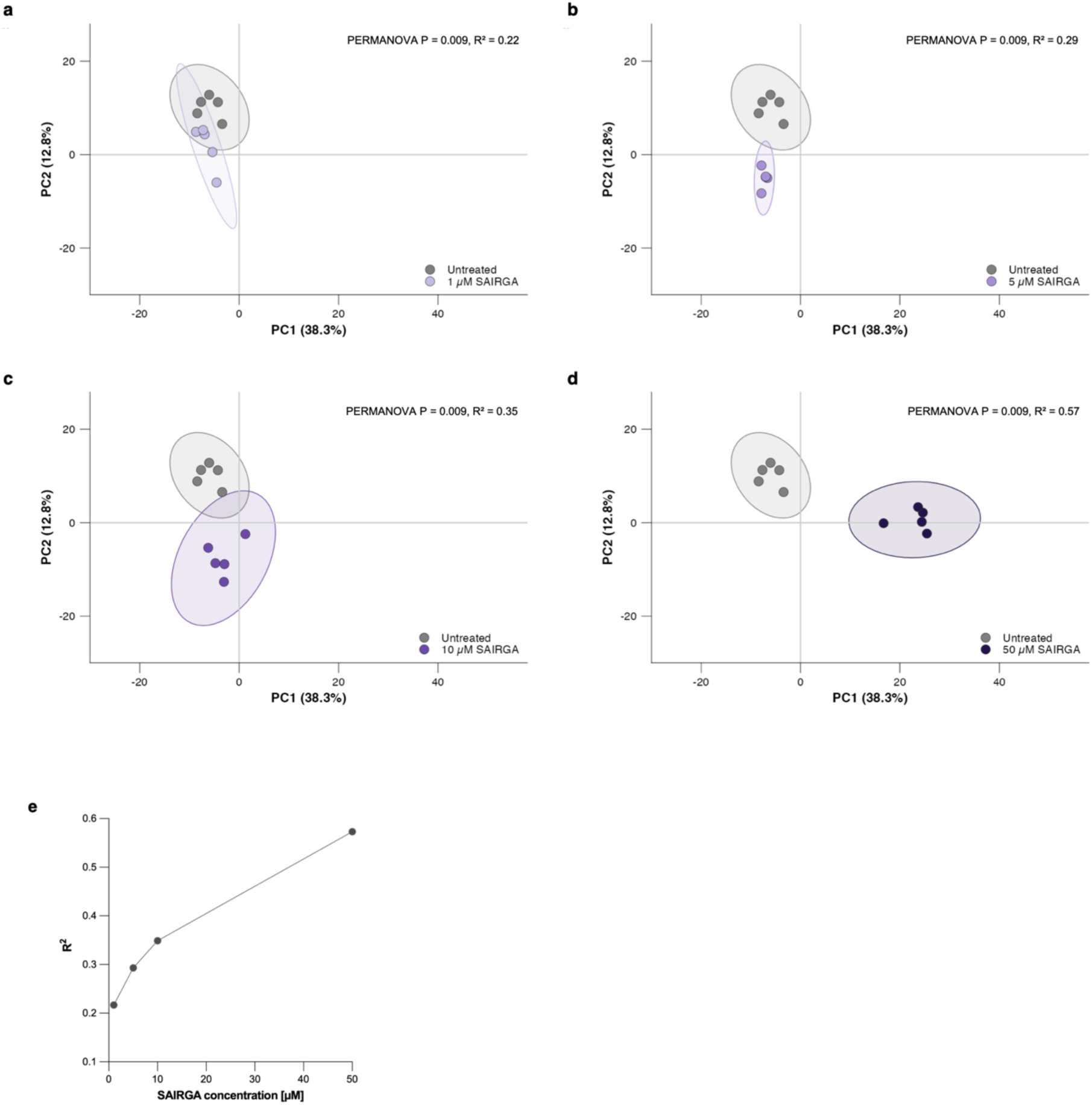
| Global metabolomic separation increases with SAIRGA concentration. **a–d**, PCA score plots comparing untreated controls with a, 1 µM, b, 5 µM, c, 10 µM and d, 50 µM SAIRGA-treated B. subtilis colony-biofilm metabolomes. Axes show PC1 (38.3%) and PC2 (12.8%) from the analysis of all 349 features. Shaded regions indicate 95% data ellipses. PERMANOVA statistics are indicated in each panel; all four comparisons were significant (P = 0.009, 999 permutations), with R² = 0.217, 0.293, 0.349 and 0.573 at 1, 5, 10 and 50 µM, respectively. e, Proportion of metabolomic variance explained by SAIRGA treatment (PERMANOVA R²) across the concentration series; increased monotonically across the concentration series. Full PERMANOVA statistics are provided in Supplementary data Table 4. n = 5 biological replicates per condition.

**Extended Data Fig. 3.**
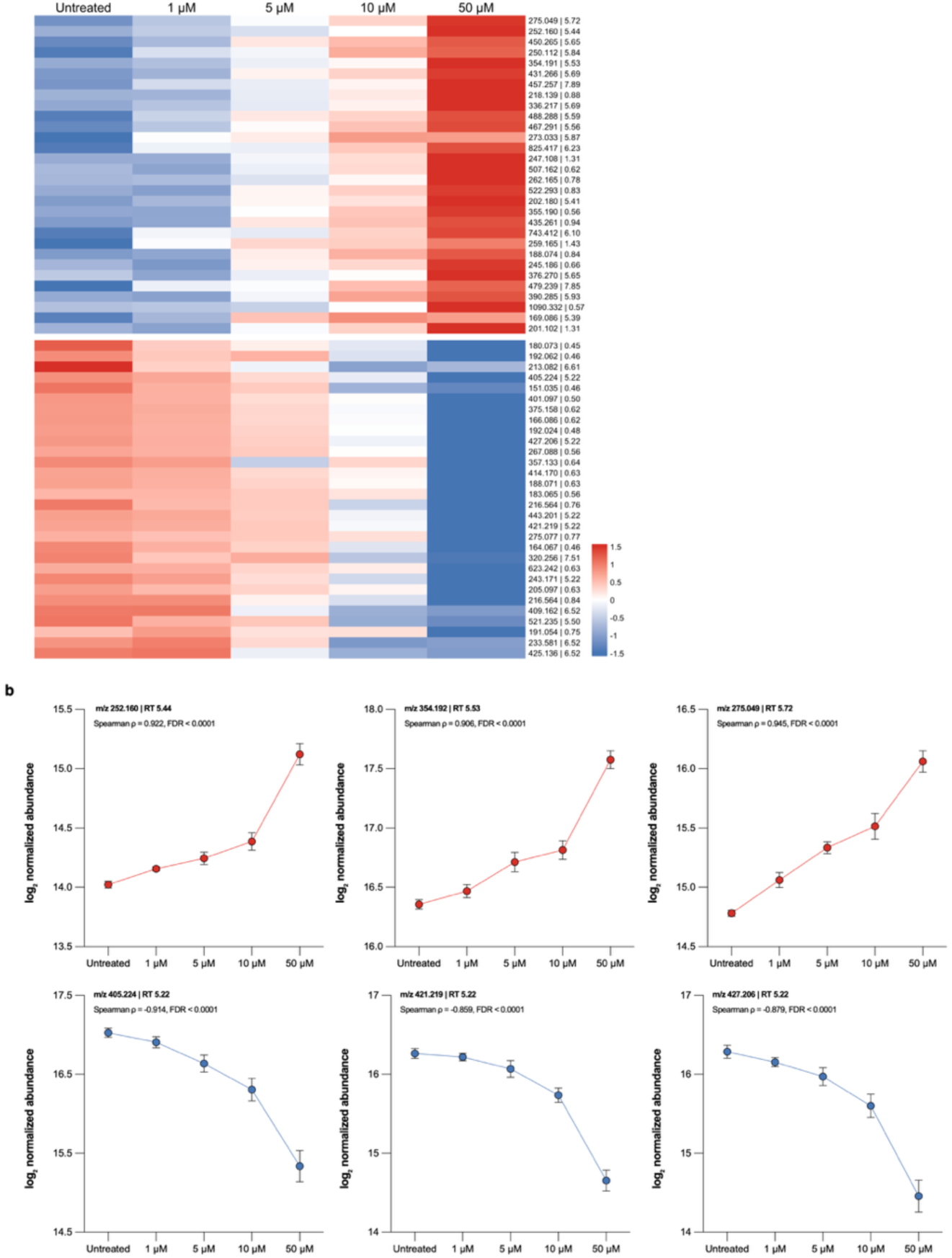
Dose-dependent metabolomic trends in SAIRGA-treated B. subtilis colony biofilms. **a**, Heatmap of the 30 most significant increasing and 30 most significant decreasing features identified by Spearman dose-response analysis (FDR < 0.05), ranked within each direction by FDR, then by |ρ| and |log₂ fold change at 50 µM|. Values are row-wise z-scores of group mean log₂-normalized abundance; columns are ordered by SAIRGA concentration. Red, above the feature mean; blue, below. Row labels indicate m/z and retention time. The complete 161-feature heatmap is shown in Extended Data Fig. 4. **b**, Dose-response profiles of six representative features: three increasing (top, red) and three decreasing (bottom, blue). All six are strictly monotonic across the concentration series, changing in the trend direction in all four consecutive dose steps. Points show group means ± s.e.m.; individual biological replicates are overlaid. Spearman ρ and Benjamini–Hochberg FDR are indicated in each panel. Feature statistics provided in Supplementary Table 6. n = 5 biological replicates per condition.

**Extended Data Fig. 4.**
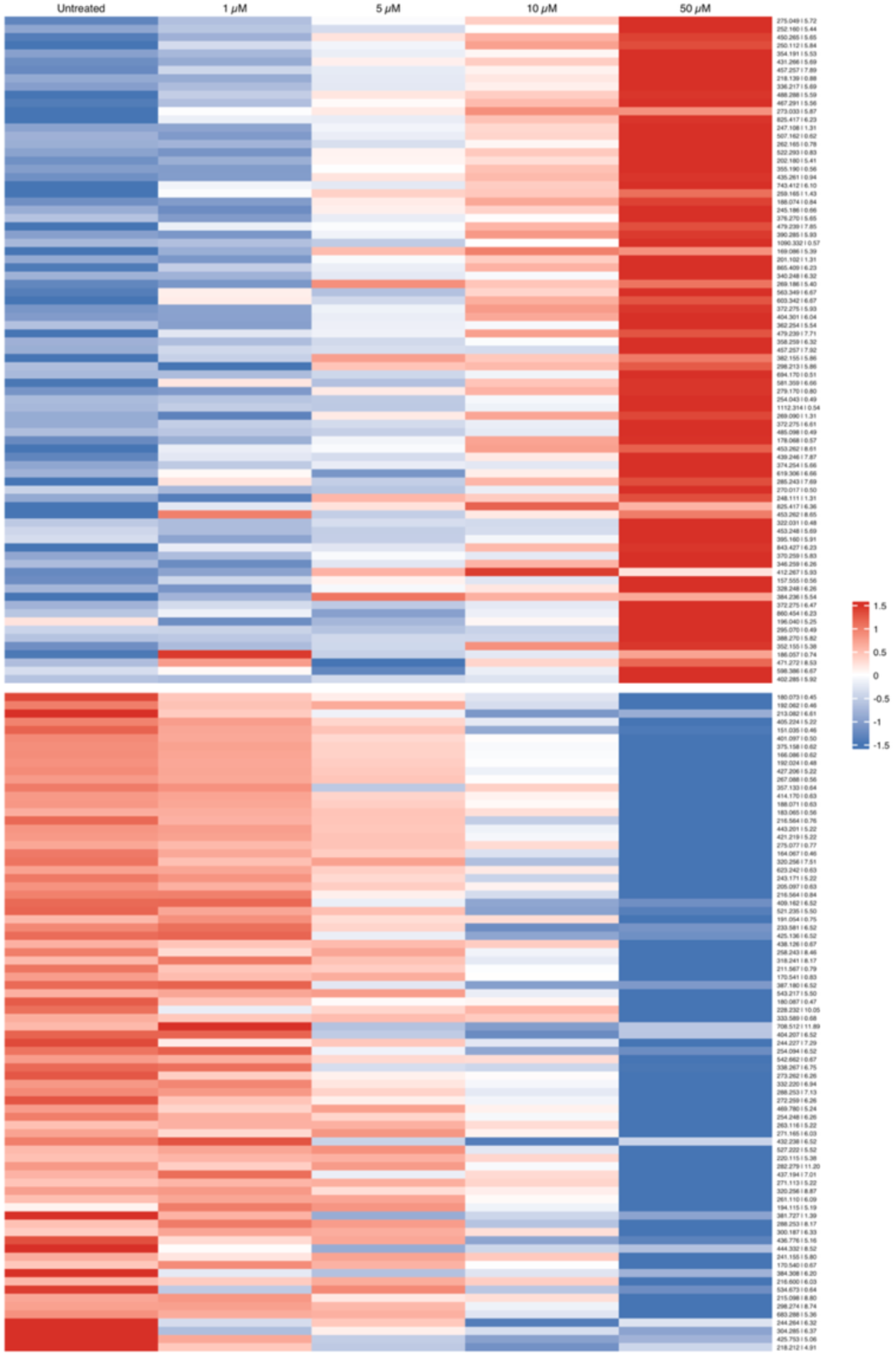
| Complete heatmap of significant SAIRGA dose-response features. Heatmap of all 161 metabolomic features with a significant monotonic dose-response trend (Spearman rank correlation, FDR < 0.05), comprising 81 increasing and 80 decreasing features. Features are ordered within each direction block by FDR, then by |rho| and |log₂ fold change at 50 µM|, using the same ranking as Extended Data Fig. 3a, which shows the top 30 of each block. Values are row-wise z-scores of group mean log₂-normalized abundance; columns are ordered by SAIRGA concentration. Red, above the feature mean; blue, below. Row labels indicate m/z and retention time. Feature statistics provided in Supplementary Table 6. n = 5 biological replicates per condition.

**Extended Data Fig. 5.**
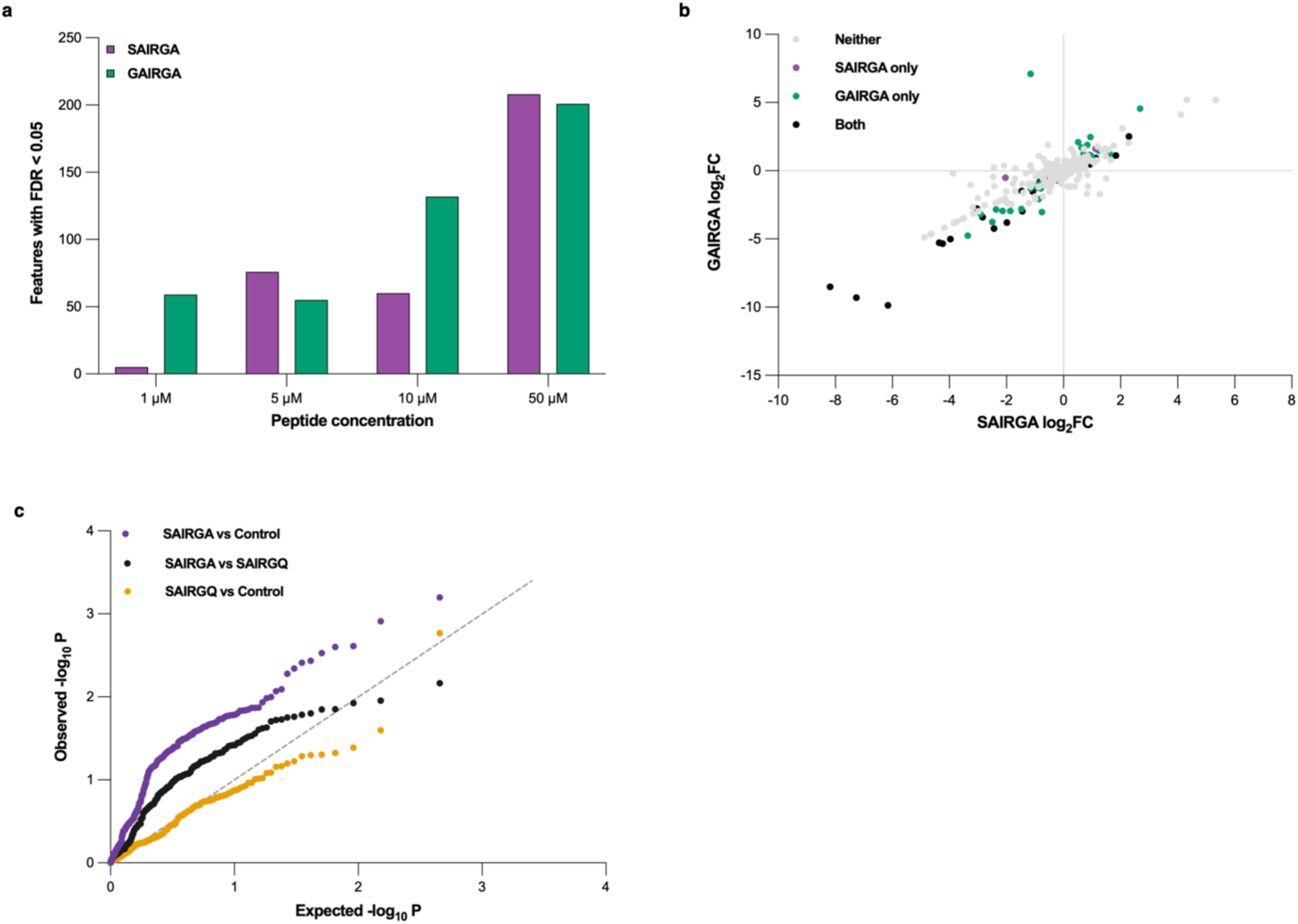
| The metabolomic response tracks peptide activity. **a**, Number of features significantly altered by SAIRGA or GAIRGA relative to untreated controls across the concentration series (Welch’s t-test with Benjamini–Hochberg correction, FDR < 0.05). SAIRGA and GAIRGA samples were processed together in a shared feature table using identical parameters and the same untreated controls. **b**, Log₂ fold changes at 10 µM across all 423 shared features. The dashed line indicates identity. Fold changes induced by SAIRGA and GAIRGA were strongly correlated (Pearson r = 0.863). n = 5 biological replicates per condition for panels **a** and **b**. **c**, Quantile–quantile plot of per-feature P values from an independent experiment comparing SAIRGA with the inactive C-terminal variant SAIRGQ and untreated controls (20 µM, negative-ionization mode). The dashed diagonal indicates the expected distribution under the null hypothesis. SAIRGA versus untreated controls yielded 97 nominally significant features (P < 0.05; empirical permutation P = 0.0018), whereas SAIRGQ versus untreated controls yielded two (empirical permutation P = 0.8108). The direct SAIRGA–SAIRGQ comparison yielded 40 nominally significant features (empirical permutation P = 0.0322). No individual feature survived Benjamini– Hochberg correction. n = 6 biological replicates per condition.

**Extended Data Fig. 6.**
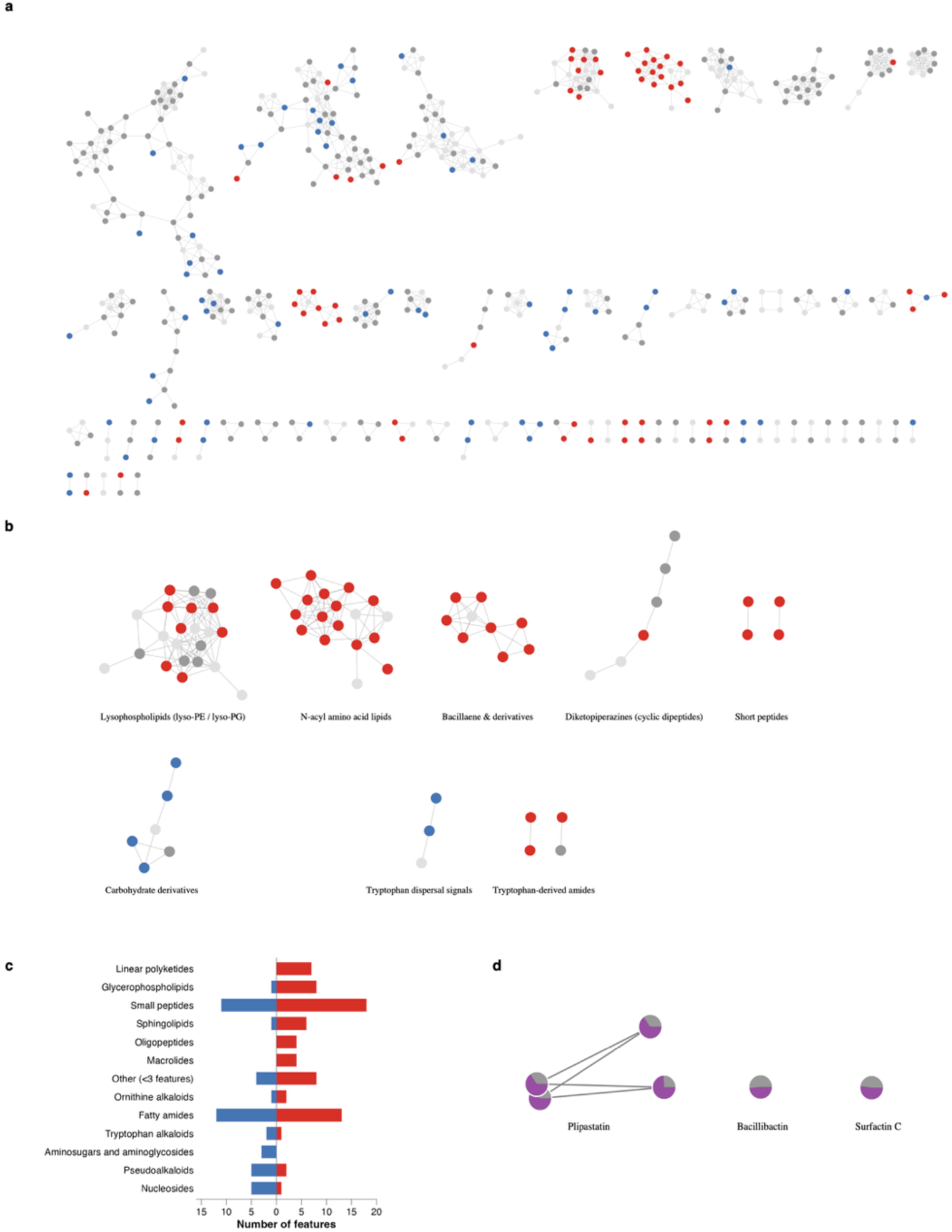
| SAIRGA reshapes the *B. subtilis* colony-biofilm metabolome across structurally diverse chemical families. **a**, Feature-based molecular network generated in GNPS2 from MS2 spectra acquired across untreated and SAIRGA- treated colony biofilms. The network comprised 605 nodes, of which 471 were connected to at least one neighboring node; 134 singletons are not shown. Node colour indicates the dose-response classification: red, significantly increasing; blue, significantly decreasing (Spearman rank correlation, Benjamini–Hochberg FDR < 0.05); dark grey, quantified but not significantly dose-responsive; light grey, not retained in the final quantitative feature set. **b**, Representative molecular families containing metabolites associated with the SAIRGA response. Families were defined by network connectivity and annotated by integrating GNPS2 spectral-library evidence with SIRIUS/CSI:FingerID structure prediction and CANOPUS chemical class assignment. Annotation confidence follows MSI levels 2–4 as defined in Methods. **c**, Distribution of significantly dose-responsive features across CANOPUS/NPClassifier chemical superclasses. Red, increasing; blue, decreasing. Of the 161 significantly trending features, 119 carried a chemical class assignment and are shown. Classes represented by fewer than three features are pooled as ‘Other’; features lacking a superclass assignment or flagged as likely contaminants are excluded. d, Relative abundance of network-detected plipastatin, bacillibactin and surfactin features at 10 µM SAIRGA relative to untreated controls. Grey, untreated; purple, SAIRGA. Bacillibactin and surfactin C occur as network singletons, whereas the two plipastatin features belong to the same molecular family. These compounds carry GNPS spectral-library annotations (MSI level 2). Their detection in the untargeted dataset, together with the broader specialized-metabolite response, motivated targeted follow-up using an acquisition method optimized for large lipopeptides (Fig. 5). n = 5 biological replicates per condition.

**Extended Data Fig. 7.**
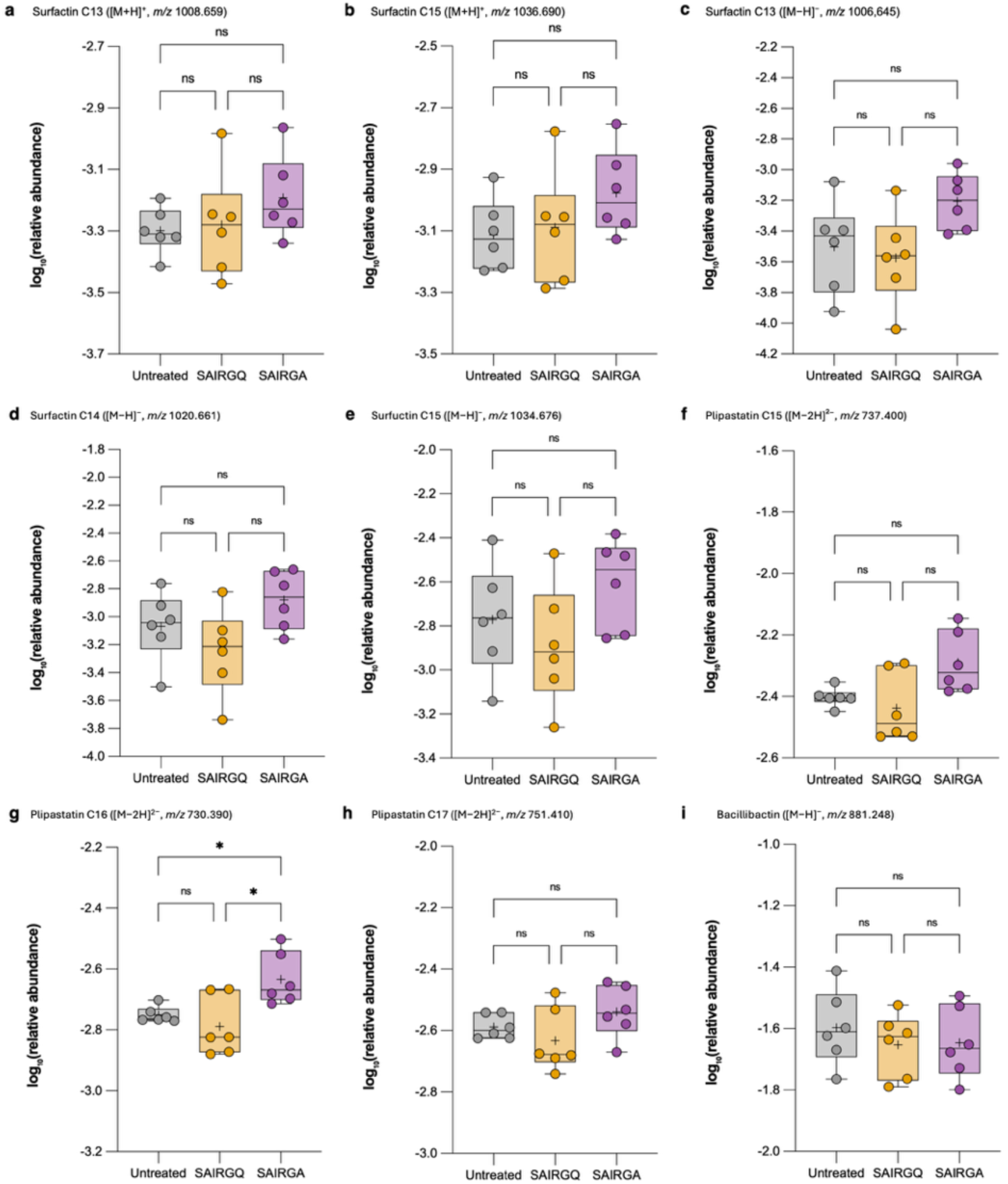
| Extended targeted profiling of *B. subtilis* specialized metabolites. Targeted LC–HRMS measurements complementing the representative metabolites shown in Fig. 5. Box plots show the median (centre line), interquartile range (box) and full data range (whiskers), with individual biological replicates overlaid; crosses indicate group means. n = 6 biological replicates per condition. Positive-ionization measurements were normalized to terfenadine and negative-ionization measurements to chlorpropamide; values are log₁₀- transformed. Statistical comparisons used Welch’s ANOVA followed by Games–Howell multiple-comparisons testing; *P < 0.05, **P < 0.01; ns, not significant. **a,b**, Surfactin C13 and C15 measured in ESI+. **c–e**, Surfactin C13, C14 and C15 measured in ESI−. **f–h**, Plipastatin C15, C16 and C17 measured in ESI−. **i**, Bacillibactin measured in ESI−. Exact *m/z*, adducts, fold changes and pairwise adjusted P values for all measurements are provided in Supplementary Table 7.

**Extended Data Fig. 8.**
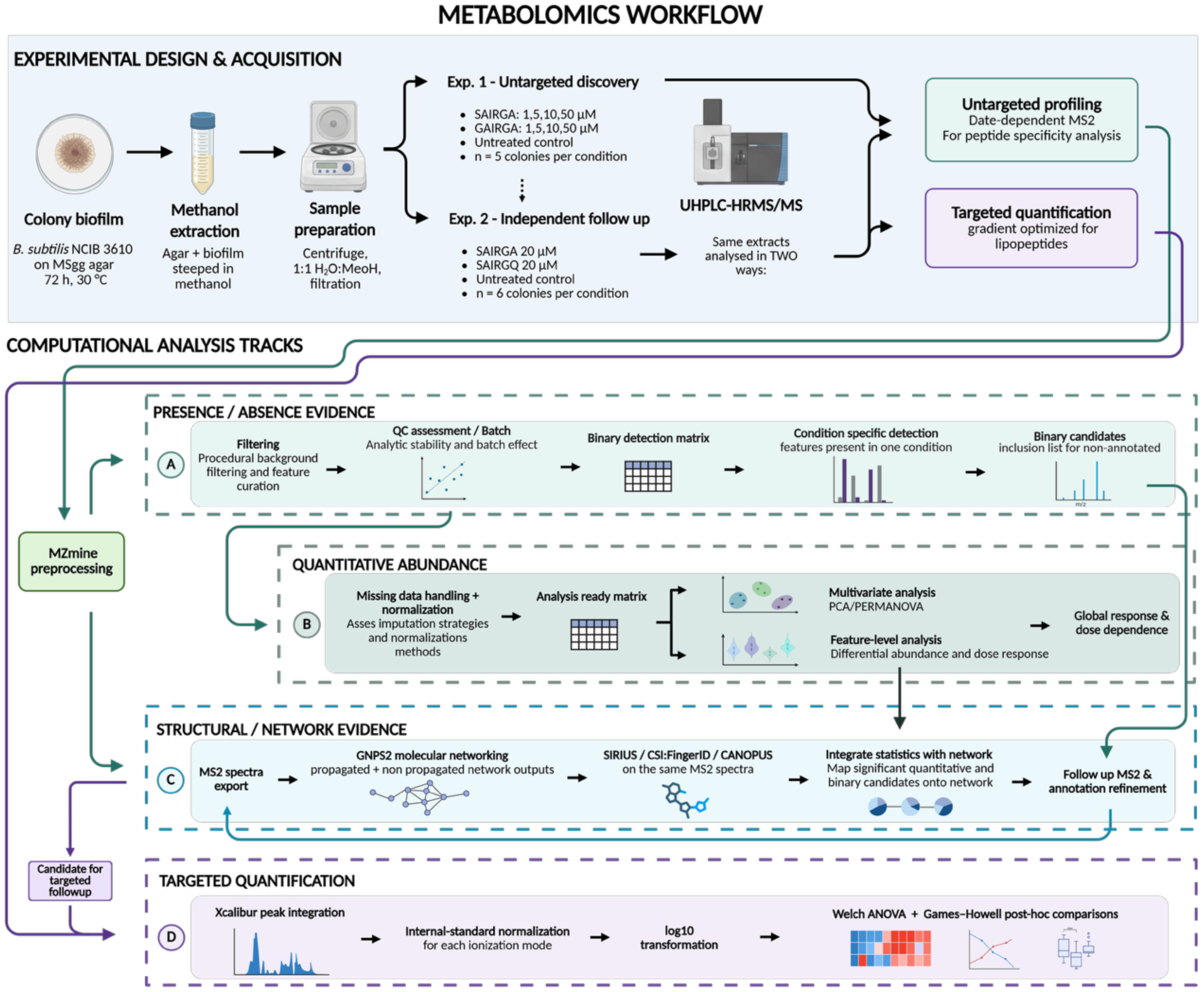
| Metabolomics experimental and analytical workflow. Overview of the experimental design and analytical strategy used to characterize the metabolic response of *Bacillus subtilis* colony biofilms to arbitrium peptides. **Experimental design and acquisition.** Experiment 1 was an untargeted discovery experiment comprising untreated colony biofilms and SAIRGA and GAIRGA concentration series (1, 5, 10 and 50 µM; n = 5 biological replicates per condition). Experiment 2 was an independent follow-up comparing untreated biofilms with SAIRGA (20 µM) and the inactive sequence variant SAIRGQ (20 µM; n = 6 per condition). The same Experiment 2 extracts were analysed in two ways: by untargeted profiling for peptide-specificity testing, and by targeted quantification using a gradient optimised for lipopeptides. **Computational analysis.** Untargeted data were preprocessed in MZmine and evaluated along complementary tracks. **(A)** Presence/absence evidence: procedural-background filtering and feature curation, assessment of analytical stability and batch structure, construction of a binary detection matrix, and identification of features detected in a condition-specific manner. Features prioritized in this track were carried forward as inclusion-list candidates for annotation, but were not used for quantitative conclusions. **(B)** Quantitative abundance: missing-value handling and normalization, followed by multivariate analysis (PCA, PERMANOVA) and feature-level analysis of differential abundance and dose response, defining the global response and its dose dependence. **(C)** Structural and network evidence: MS2 spectra were exported for feature-based molecular networking in GNPS2 and for complementary annotation with SIRIUS/CSI:FingerID and CANOPUS on the same spectra. Statistical results from tracks A and B were mapped onto the network, and selected stored extracts were re-analyzed with inclusion lists to obtain additional MS2 spectra for prioritized features; these follow-up acquisitions supported annotation only and contributed no quantitative measurements. Refined annotations were fed back into the networking step in an iterative cycle. **(D)** Targeted quantification: peak integration in Xcalibur, internal-standard normalization within each ionization mode, log₁₀ transformation, and comparison by Welch’s ANOVA with Games–Howell post-hoc testing. Positive-mode targeted measurements were normalized to terfenadine; the extracts were subsequently re-injected with chlorpropamide added for negative-mode normalization. The workflow provided four complementary outputs — global dose-dependent response, molecular-family and chemical-class annotation, independent peptide-specificity testing, and targeted validation of specialized metabolites — supporting the analyses in Figs. 4 and 5 and Extended Data Figs. 2–7.

**Extended Data Fig. 9.**
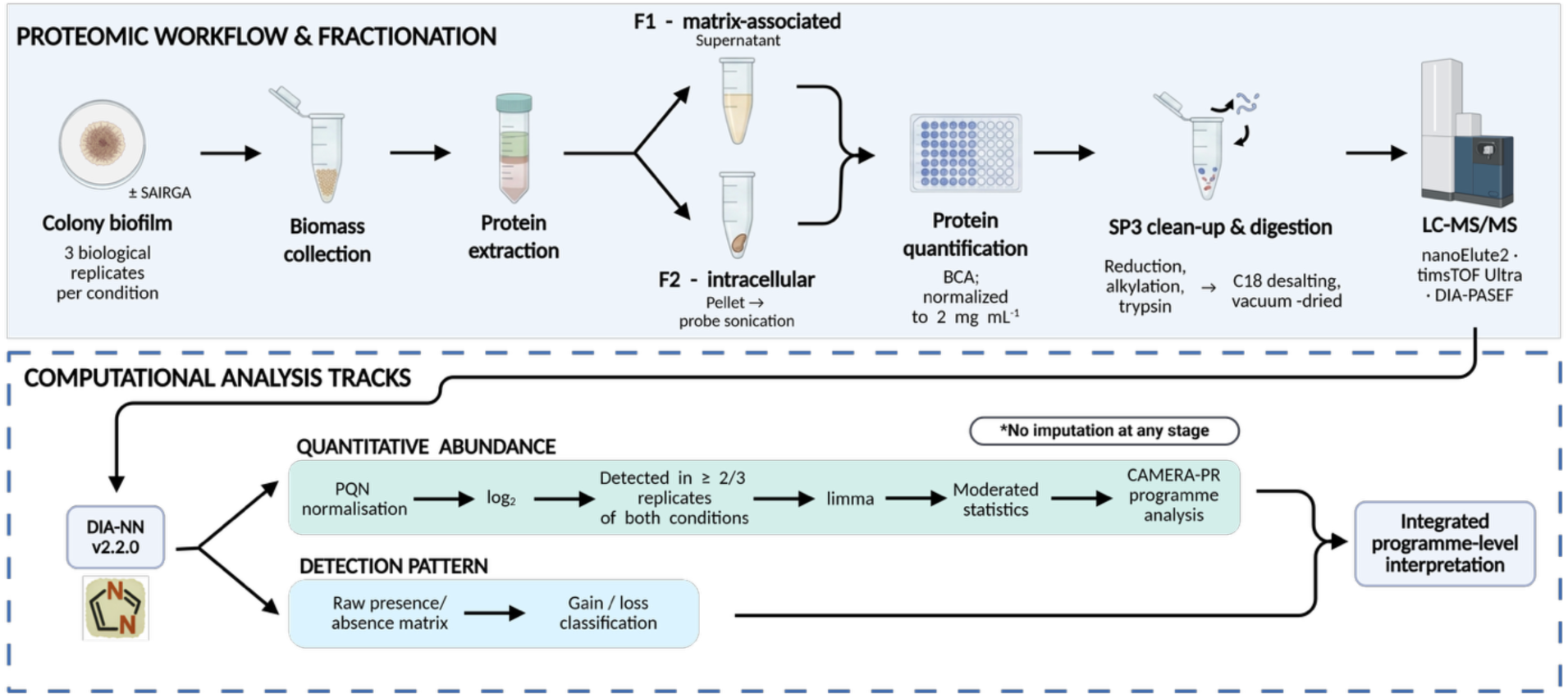
| Proteomic extraction and analysis workflow. Colony biofilms grown with or without SAIRGA were harvested and fractionated by centrifugation into a matrix-associated/extracellular-enriched fraction (F1) and an intracellular/cell-associated-enriched fraction (F2). Both fractions underwent SP3 clean-up, tryptic digestion and LC–MS/MS analysis by DIA-PASEF, followed by protein-group assignment with DIA-NN. Each fraction was analyzed along two parallel tracks. In the **quantitative-abundance track**, protein intensities were probabilistic-quotient-normalized, log₂-transformed and restricted to proteins detected in at least two of three biological replicates in both conditions. Differential abundance was analyzed with limma, and moderated *t*-statistics were used for CAMERA-PR program analysis. In the **detection-pattern track**, the untransformed presence/absence matrix was examined directly and proteins were classified using predefined detection-gain and detection-loss patterns. Missing values were not imputed in either track, and proteins represented only in the detection-pattern analysis therefore have no quantitative fold change. The two analytical tracks were combined only at the level of biological interpretation.

**Extended Data Fig. 10.**
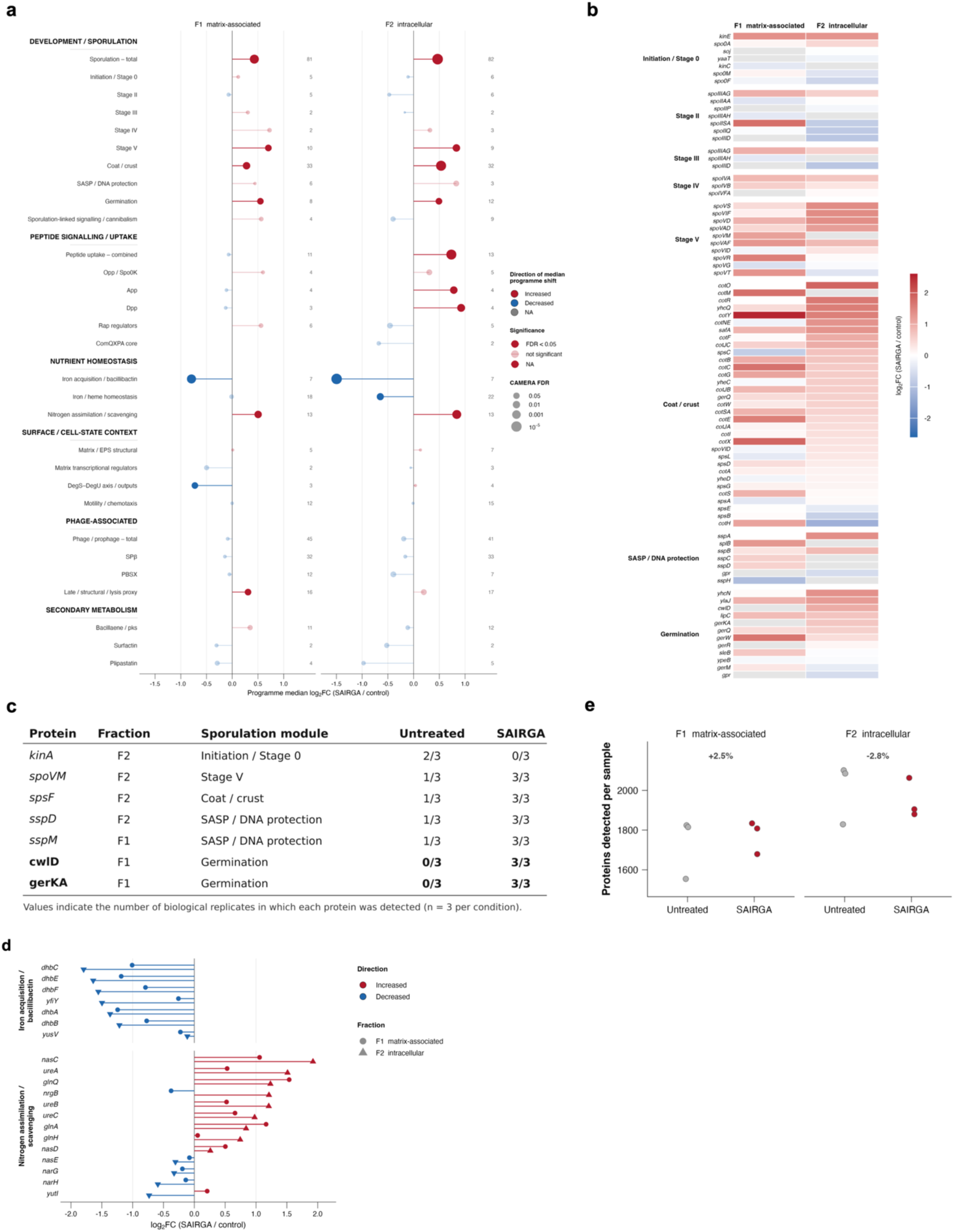
| Protein-level structure and detection context of the SAIRGA proteomic response. **a**, Complete CAMERA-PR landscape of all testable curated functional programs in F1 and F2. Point position indicates program median log₂ fold change (SAIRGA versus untreated), color indicates direction, point size reflects CAMERA-PR significance, and saturated symbols denote Benjamini–Hochberg-adjusted FDR < 0.05. Numbers indicate the number of detected proteins contributing to each program. The ComQXPA core program was not testable in F1 and is indicated as n.t. b. Protein-level log₂ fold changes for sporulation- associated proteins grouped by developmental stage and shown separately for F1 and F2. Red and blue indicate higher and lower abundance, respectively, in SAIRGA-treated biofilms; grey indicates unavailable quantitative values. Proteins assigned to more than one curated module are shown in each applicable module. **c**, Descriptive detection patterns among sporulation-associated proteins. Values indicate the number of biological replicates in which each protein was detected in untreated and SAIRGA-treated samples (n = 3 per condition). GerKA and CwlD were undetected in all untreated F1 samples and detected in all three SAIRGA-treated F1 samples. Detection changes are shown descriptively and are not treated as independent inferential evidence. d, Protein-level log₂ fold changes for CAMERA-PR members of the iron-acquisition/bacillibactin and nitrogen-assimilation/scavenging programs. Circles denote F1 and triangles F2; red and blue indicate increased and decreased abundance, respectively. e, Per-sample protein-detection depth in F1 and F2. Points represent individual biological samples. SAIRGA-treated samples contained approximately 2.5% more detected proteins in F1 and 2.8% fewer in F2 than untreated samples, providing background context for interpretation of the descriptive detection-gain/loss analysis.

## Notes

### Competing Interest Statement

The authors have declared no competing interest.

### Summary of Updates

We updated the acknowledgements section. No additional changes were made in this revision.

### Funding

- Israel Science Foundation: 2015/24

