## Supplemental Figures for "A phage communication peptide alters *Bacillus subtilis* colony development and promotes sporulation"

### Supplementary Information

**Supplementary Fig. 1 | Chemical structures and ESI-HRMS characterization of synthetic peptides.**

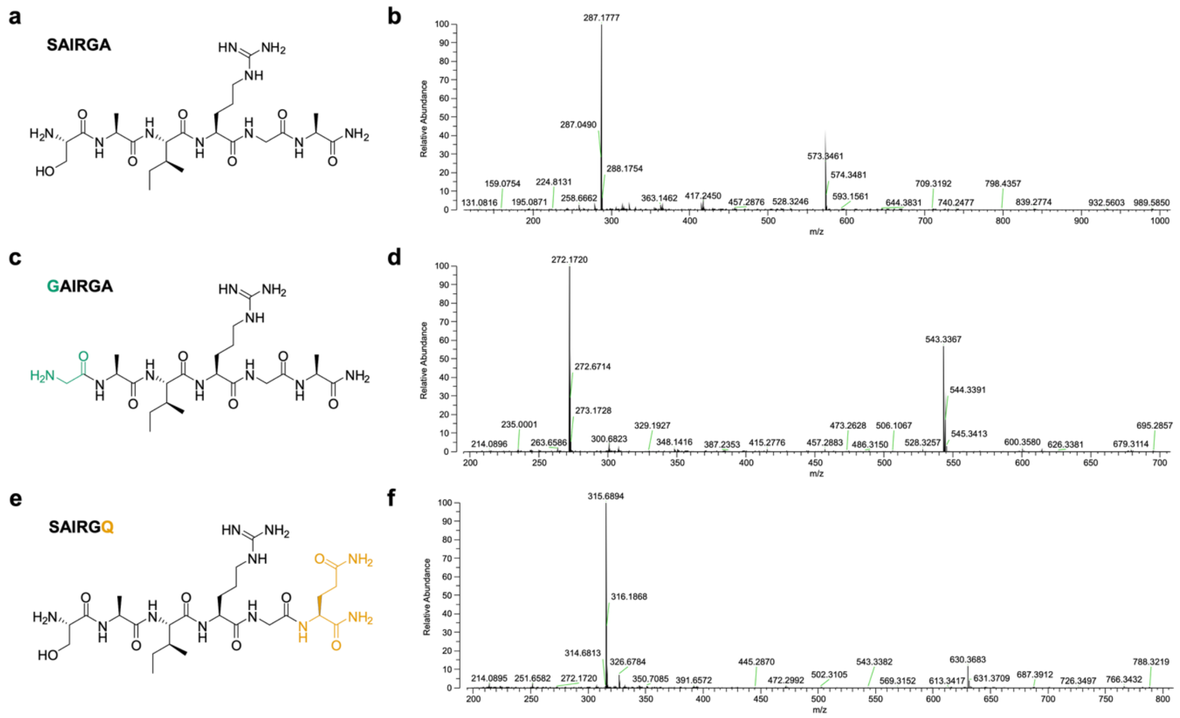


**a,c,e,** Chemical structures of SAIRGA, GAIRGA and SAIRGQ. Amino acid substitutions relative to SAIRGA are highlighted in green for GAIRGA and amber for SAIRGQ. **b,d,f,** Positive-ion electrospray ionization high-resolution mass spectra of the purified peptides. SAIRGA: calculated *m/z* 573.3467 for $[M+H]^+$ and 287.1770 for $[M+2H]^{2+}$, found 573.35 and 287.18; GAIRGA: calculated *m/z* 543.3362 for $[M+H]^+$ and 272.1717 for $[M+2H]^{2+}$, found 543.34 and 272.17; SAIRGQ: calculated *m/z* 630.3682 for $[M+H]^+$ and 315.6877 for $[M+2H]^{2+}$, found 630.37 and 315.69.

**Supplementary Table 1 | Morphometric analysis of colony biofilms.**

| Condition | Area (mm²) | Perimeter (mm) | Feret (mm) | MinFeret (mm) | Circularity | Solidity | AR | Roundness |
|---|---|---|---|---|---|---|---|---|
| **Untreated** | 377 ± 74 | 79.6 ± 9.1 | 23.7 ± 2.8 | 21.7 ± 2.3 | 0.743 ± 0.029 | 0.948 ± 0.005 | 1.03 ± 0.01 | 0.971 ± 0.007 |
| **SAIRGA 5 µM** | 1035 ± 179 | 121.5 ± 10.4 | 39.5 ± 4.3 | 35.1 ± 2.9 | 0.876 ± 0.006 | 0.971 ± 0.005 | 1.08 ± 0.04 | 0.928 ± 0.037 |
| **SAIRGA 20 µM** | 2500 ± 172 | 181.3 ± 5.7 | 58.7 ± 2.0 | 55.1 ± 2.6 | 0.954 ± 0.009 | 0.991 ± 0.004 | 1.03 ± 0.01 | 0.967 ± 0.011 |
| **GAIRGA 20 µM** | 2503 ± 133 | 181.6 ± 3.9 | 58.2 ± 1.1 | 55.5 ± 1.8 | 0.953 ± 0.012 | 0.989 ± 0.001 | 1.03 ± 0.01 | 0.972 ± 0.008 |
| **SAIRGQ 20 µM** | 388 ± 58 | 80.0 ± 5.4 | 24.0 ± 1.6 | 21.9 ± 1.1 | 0.765 ± 0.119 | 0.952 ± 0.023 | 1.04 ± 0.03 | 0.961 ± 0.031 |

Values are mean ± SD; n = 3 biological replicates per group. Colony boundaries were manually delineated in ImageJ/Fiji using the outermost continuous bacterial biomass, after spatial calibration to the internal plate diameter (60 mm). Area and diameters (Feret, MinFeret) describe colony size/spreading; Circularity ($4\pi \cdot \text{Area}/\text{Perimeter}^2$), Solidity, aspect ratio (AR) and Roundness describe colony shape (values near 1 indicate a smooth, round outline). Area was the primary readout (Fig. 1c); perimeter-dependent parameters are more sensitive to manual delineation and are reported for completeness.

**Supplementary Fig. 2 | SAIRGA does not produce an obvious macroscopic change in static-liquid pellicles under the conditions tested**

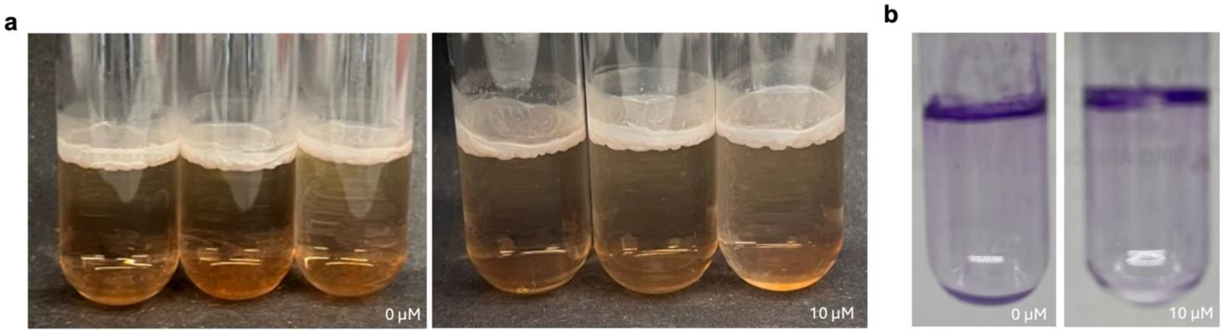


*B. subtilis* NCIB 3610 was grown under static conditions in MSgg medium in the absence or presence of 10 µM SAIRGA for 72 h at 30 °C. **a,** Side-view images of three independent biological replicates showing pellicle and ring formation at the air–liquid–wall interface. **b,** Representative crystal-violet-stained tubes following removal of the culture medium and washing. No obvious difference in pellicle- or ring-associated staining was observed between untreated and SAIRGA-treated cultures under the conditions tested. Crystal-violet staining was assessed qualitatively and was not quantified spectrophotometrically.

**Supplementary Fig. 3 | Chemical structures and ESI-HRMS characterization of GMPRGA and D-SAIRGA.**

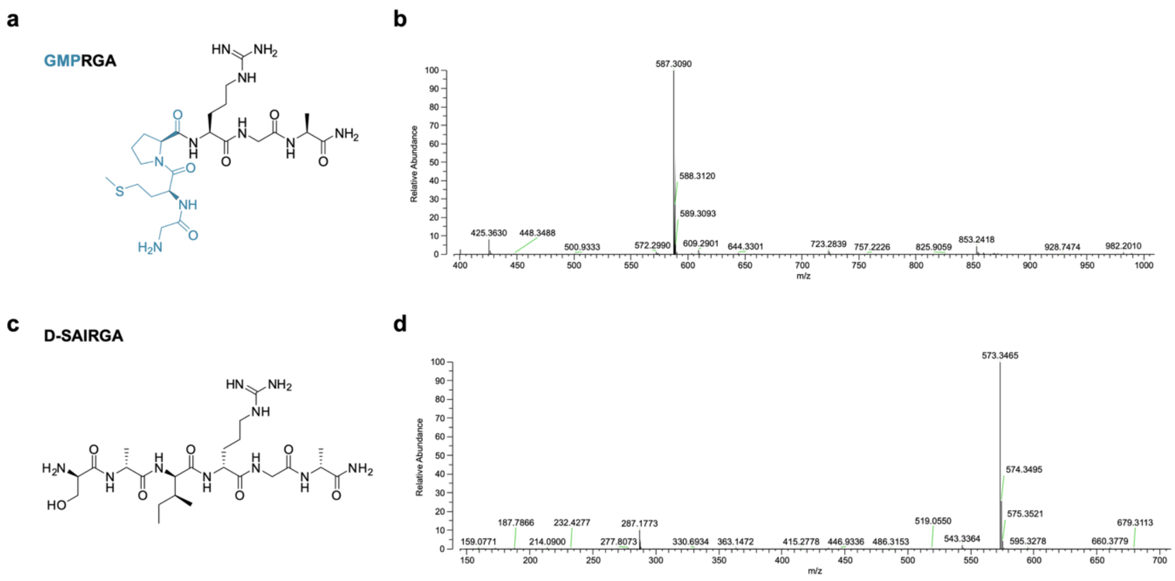


**a,c,** Chemical structures of the C-terminally amidated synthetic peptides GMPRGA and D-SAIRGA, respectively. **b,d,**Positive-ion electrospray ionization high-resolution mass spectra of the purified peptides. GMPRGA: calculated *m/z*587.3082 for $[M+H]^+$, found 587.3090; D-SAIRGA: calculated *m/z* 573.3467 for $[M+H]^+$, found 573.3465. D-SAIRGA is the all-D enantiomer of SAIRGA and therefore has the same elemental composition and exact mass as L-SAIRGA.

**Supplementary Table 2 | Morphometric quantification of colony biofilms used to assess peptide sequence, stereochemistry, and AimR dependence.**

| Condition | Area (mm²) | Perimeter (mm) | Feret (mm) | MinFeret (mm) | Circularity | Solidity | AR | Roundness |
|---|---|---|---|---|---|---|---|---|
| WT - Untreated | 391 ± 79 | 77.8 ± 5.6 | 23.5 ± 2.0 | 22.1 ± 2.3 | 0.805 ± 0.063 | 0.971 ± 0.011 | 1.018 ± 0.012 | 0.982 ± 0.012 |
| WT- SAIRGA 20 μM | 1018 ± 172 | 119.0 ± 9.1 | 37.6 ± 3.2 | 35.2 ± 2.9 | 0.899 ± 0.022 | 0.981 ± 0.004 | 1.039 ± 0.012 | 0.963 ± 0.011 |
| WT - GMPRGA 20 μM | 1546 ± 302 | 145.2 ± 16.0 | 46.0 ± 4.9 | 43.4 ± 4.1 | 0.917 ± 0.019 | 0.984 ± 0.001 | 1.037 ± 0.025 | 0.965 ± 0.023 |
| WT - GAIRGA 20 μM | 1231 ± 85 | 131.8 ± 7.7 | 41.8 ± 1.3 | 38.6 ± 1.5 | 0.892 ± 0.043 | 0.983 ± 0.005 | 1.044 ± 0.020 | 0.958 ± 0.018 |
| WT - Untreated | 421 ± 45 | 76.8 ± 3.6 | 24.3 ± 1.5 | 22.9 ± 1.0 | 0.894 ± 0.013 | 0.978 ± 0.005 | 1.034 ± 0.018 | 0.967 ± 0.017 |
| WT - SAIRGA 20 μM | 857 ± 88 | 109.2 ± 4.0 | 34.8 ± 0.8 | 31.7 ± 2.8 | 0.901 ± 0.029 | 0.983 ± 0.004 | 1.077 ± 0.091 | 0.933 ± 0.076 |
| WT - D-SAIRGA 20 μM | 513 ± 110 | 84.7 ± 9.7 | 26.9 ± 3.1 | 25.0 ± 2.7 | 0.893 ± 0.012 | 0.982 ± 0.002 | 1.039 ± 0.027 | 0.963 ± 0.025 |
| **ΔaimR -** Untreated | 630 ± 42 | 100.2 ± 2.9 | 31.0 ± 1.2 | 27.7 ± 0.6 | 0.792 ± 0.096 | 0.958 ± 0.015 | 1.068 ± 0.062 | 0.938 ± 0.052 |
| **ΔaimR -** SAIRGA 20 μM | 2513 ± 248 | 181.8 ± 9.5 | 58.9 ± 2.2 | 55.4 ± 3.2 | 0.954 ± 0.009 | 0.991 ± 0.003 | 1.032 ± 0.022 | 0.970 ± 0.021 |

Values are mean ± s.d.; n = 3 independent biological replicates per group. Panels a–c correspond to separate biological experiments and therefore include separate contemporaneous controls. Colony boundaries were manually delineated in ImageJ/Fiji using the outermost continuous bacterial biomass after spatial calibration to the internal plate diameter (60 mm). Area and diameters (Feret, MinFeret) describe colony size/spreading; Circularity ($4\pi \cdot \text{Area}/\text{Perimeter}^2$), Solidity, aspect ratio (AR) and Roundness describe colony shape. Area was the primary readout in Fig. 2; perimeter-dependent parameters are more sensitive to manual delineation and are reported for completeness.

**Supplementary Table 3 | Feature filtering cascade for untargeted metabolomic analysis**

Primary quantitative feature-filtering cascade for Experiment 1. Feature detection used a minimum peak-height threshold of $5 \times 10^5$. MS2 acquisition and molecular-networking counts were tracked separately from this quantitative filtering cascade.

| Processing stage | Features | Notes |
|---|---|---|
| MZmine feature detection | 606 | *Minimun peak height 5x10$^5$* |
| After blank subtraction | 434 | *Features retained above the procedural blanks* |
| **After artifact deduplication** | **349** | **82 splitting artifacts, 2 all-zero features and 1 chromatographic artifact removed** |

Chromatographic splitting artifacts were identified as features sharing accurate mass (±0.005 Da) in contiguous retention-time ladders that collapsed under independent re-integration with revised peak-resolution parameters; series reproduced by re-integration were retained as genuine isomers. Two features with no detections in any sample and one shown by targeted re-injection to be a chromatographic artifact were also removed. Per-feature decisions are given in the source data.

**Supplementary Table 4 | PERMANOVA statistics for global metabolomic separation.**

Permutational multivariate analysis of variance (adonis2, vegan package) on Euclidean distances of the normalised $\log_2$ matrix (349 features × 25 samples), 999 permutations. $R^2$, proportion of total metabolomic variance explained by treatment group. n = 5 independently grown biofilm colonies per condition.

| Comparison | pseudo-F | $R^2$ | Variance explained | P value |
|---|---|---|---|---|
| Untreated vs 1 μM | 2.21 | 0.217 | 21.7% | 0.009 |
| Untreated vs 5 μM | 3.32 | 0.293 | 29.3% | 0.009 |
| Untreated vs 10 μM | 4.29 | 0.349 | 34.9% | 0.009 |
| Untreated vs 50 μM | 10.72 | 0.573 | 57.3% | 0.009 |
| **Global (all five groups)** | **5.82** | **0.538** | **53.8%** | **0.001** |

$R^2$ increases monotonically with SAIRGA concentration (Spearman rho = 1.00), indicating that the magnitude of the global metabolomic shift, and not only the number of individual features affected, is dose-dependent

**Supplementary Table 5 | Differential abundance of metabolomic features at each SAIRGA concentration.**

Features significantly altered relative to untreated controls (Welch's t-test with Benjamini–Hochberg correction), from a set of 349 features. Up and Down indicate the direction of the $\log_2$ fold change. Nominal P < 0.05 is shown for reference against the 17 features expected by chance. Light-grey values denote comparisons with no feature surviving FDR correction. n = 5 biological replicates per condition. n = 5 biological replicates per condition.

| Concentration | FDR < 0.05 | Up | Down | Nominal P < 0.05 | Minimum FDR |
|---|---|---|---|---|---|
| 1 µM | 0 | 0 | 0 | 74 | 0.051 |
| 5 µM | **88** | 55 | 33 | 128 | 0.0016 |
| 10 µM | **103** | 62 | 41 | 155 | $< 10^{-4}$ |
| 50 µM | **193** | 98 | 95 | 219 | $< 10^{-4}$ |

At 1 µM, no individual feature survived FDR correction, although the metabolome as a whole was already significantly separated from untreated controls (PERMANOVA P = 0.009; Supplementary Table 4). Feature filtering is described in Supplementary Table 3; per-feature statistics are provided in the source data.

**Supplementary Table 6 | Metabolomic features with significant SAIRGA dose-response trends.**

All 161 features with a significant monotonic trend across SAIRGA concentrations (Spearman rank correlation, FDR < 0.05). Steps, number of consecutive dose steps (of four) in which the feature moved in the direction of its overall trend. Annotations, where available, are provided in the source data. n = 5 biological replicates per condition.

**Increasing with dose (n = 81)**

| m/z | RT (min) | rho | FDR | Steps |
|---|---|---|---|---|
| 275.0486 | 5.72 | 0.945 | 1.96e-10 | 4 |
| 252.1597 | 5.44 | 0.922 | 6.12e-09 | 4 |
| 450.2646 | 5.65 | 0.918 | 6.12e-09 | 4 |
| 250.1122 | 5.84 | 0.914 | 7.7e-09 | 4 |
| 354.1915 | 5.53 | 0.906 | 1.61e-08 | 4 |
| 431.2659 | 5.69 | 0.902 | 2.3e-08 | 4 |
| 218.1389 | 0.88 | 0.894 | 3.94e-08 | 4 |
| 457.2567 | 7.89 | 0.894 | 3.94e-08 | 4 |
| 336.2173 | 5.69 | 0.883 | 9.86e-08 | 4 |
| 488.2877 | 5.59 | 0.883 | 9.86e-08 | 4 |
| 467.2913 | 5.56 | 0.871 | 2.2e-07 | 4 |
| 247.1079 | 1.31 | 0.867 | 2.61e-07 | 4 |
| 273.0329 | 5.87 | 0.867 | 2.61e-07 | 3 |
| 825.4170 | 6.23 | 0.867 | 2.61e-07 | 3 |
| 507.1624 | 0.62 | 0.863 | 3.45e-07 | 3 |
| 262.1651 | 0.78 | 0.851 | 6.74e-07 | 4 |
| 522.2930 | 0.83 | 0.851 | 6.74e-07 | 3 |
| 202.1804 | 5.41 | 0.839 | 1.37e-06 | 4 |
| 355.1898 | 0.56 | 0.835 | 1.61e-06 | 3 |
| 435.2607 | 0.94 | 0.835 | 1.61e-06 | 4 |

| m/z | RT (min) | rho | FDR | Steps |
|---|---|---|---|---|
| 188.0742 | 0.84 | 0.828 | 2.47e-06 | 4 |
| 259.1654 | 1.43 | 0.828 | 2.47e-06 | 4 |
| 743.4118 | 6.10 | 0.828 | 2.47e-06 | 3 |
| 245.1861 | 0.66 | 0.824 | 2.95e-06 | 3 |
| 376.2699 | 5.65 | 0.824 | 2.95e-06 | 3 |
| 479.2388 | 7.85 | 0.820 | 3.65e-06 | 4 |
| 390.2852 | 5.93 | 0.796 | 1.27e-05 | 3 |
| 1090.3318 | 0.57 | 0.796 | 1.27e-05 | 4 |
| 169.0861 | 5.39 | 0.792 | 1.5e-05 | 3 |
| 201.1024 | 1.31 | 0.792 | 1.5e-05 | 3 |
| 865.4093 | 6.23 | 0.784 | 2.16e-05 | 4 |
| 340.2484 | 6.32 | 0.781 | 2.55e-05 | 4 |
| 269.1861 | 5.40 | 0.773 | 3.32e-05 | 3 |
| 362.2541 | 5.54 | 0.773 | 3.32e-05 | 3 |
| 372.2747 | 5.93 | 0.773 | 3.32e-05 | 4 |
| 404.3009 | 6.04 | 0.773 | 3.32e-05 | 4 |
| 563.3486 | 6.67 | 0.773 | 3.32e-05 | 3 |
| 603.3415 | 6.67 | 0.773 | 3.32e-05 | 3 |
| 358.2588 | 6.32 | 0.769 | 3.78e-05 | 4 |
| 479.2388 | 7.71 | 0.769 | 3.78e-05 | 4 |
| 457.2567 | 7.92 | 0.765 | 4.36e-05 | 3 |
| 382.1546 | 5.86 | 0.757 | 5.92e-05 | 3 |
| 298.2127 | 5.86 | 0.749 | 7.94e-05 | 3 |
| 694.1700 | 0.51 | 0.745 | 9.17e-05 | 4 |
| 581.3593 | 6.66 | 0.740 | 0.000112 | 3 |
| 279.1704 | 0.80 | 0.737 | 0.00012 | 4 |
| 254.0432 | 0.49 | 0.733 | 0.000137 | 4 |
| 1112.3144 | 0.54 | 0.730 | 0.000157 | 3 |
| 269.0899 | 1.31 | 0.726 | 0.000174 | 3 |
| 372.2747 | 6.61 | 0.714 | 0.000255 | 4 |
| 485.0978 | 0.49 | 0.714 | 0.000255 | 4 |
| 178.0680 | 0.57 | 0.702 | 0.000367 | 4 |
| 453.2617 | 8.61 | 0.698 | 0.000408 | 4 |
| 439.2459 | 7.87 | 0.679 | 0.000731 | 3 |
| 374.2542 | 5.66 | 0.663 | 0.00109 | 3 |
| 285.2425 | 7.69 | 0.659 | 0.00119 | 3 |
| 619.3060 | 6.66 | 0.659 | 0.00119 | 3 |
| 270.0172 | 0.50 | 0.647 | 0.00158 | 3 |
| 248.1112 | 1.31 | 0.628 | 0.00247 | 2 |
| 825.4171 | 6.36 | 0.624 | 0.0027 | 3 |
| 453.2617 | 8.65 | 0.620 | 0.00292 | 3 |
| 322.0309 | 0.48 | 0.612 | 0.00344 | 2 |
| 453.2478 | 5.69 | 0.588 | 0.00566 | 3 |
| 395.1605 | 5.91 | 0.584 | 0.00597 | 3 |
| 843.4275 | 6.23 | 0.584 | 0.00597 | 3 |
| 346.2589 | 6.26 | 0.581 | 0.0064 | 3 |
| 370.2591 | 5.83 | 0.581 | 0.0064 | 3 |
| 412.2673 | 5.93 | 0.577 | 0.00685 | 3 |
| 157.5547 | 0.56 | 0.573 | 0.00739 | 3 |
| 328.2483 | 6.26 | 0.553 | 0.0109 | 3 |
| 384.2360 | 5.54 | 0.553 | 0.0109 | 3 |
| 372.2747 | 6.47 | 0.545 | 0.0125 | 3 |
| 860.4543 | 6.23 | 0.518 | 0.0196 | 3 |
| 196.0397 | 5.25 | 0.506 | 0.0236 | 3 |
| 295.0699 | 0.49 | 0.482 | 0.0335 | 2 |
| 352.1547 | 5.38 | 0.479 | 0.0352 | 4 |
| 388.2696 | 5.82 | 0.479 | 0.0352 | 3 |
| 186.0568 | 0.74 | 0.471 | 0.0393 | 3 |

| m/z | RT (min) | rho | FDR | Steps |
|---|---|---|---|---|
| 471.2724 | 8.53 | 0.459 | 0.0462 | 3 |
| 402.2853 | 5.92 | 0.455 | 0.0483 | 3 |
| 598.3855 | 6.67 | 0.455 | 0.0483 | 3 |

**Decreasing with dose (n = 80)**

| m/z | RT (min) | rho | FDR | Steps |
|---|---|---|---|---|
| 180.0731 | 0.45 | -0.981 | 3.12e-15 | 4 |
| 192.0619 | 0.46 | -0.918 | 6.12e-09 | 3 |
| 213.0824 | 6.61 | -0.918 | 6.12e-09 | 3 |
| 405.2239 | 5.22 | -0.914 | 7.7e-09 | 4 |
| 151.0354 | 0.46 | -0.906 | 1.61e-08 | 4 |
| 375.1583 | 0.62 | -0.898 | 3e-08 | 4 |
| 401.0966 | 0.50 | -0.898 | 3e-08 | 4 |
| 166.0864 | 0.62 | -0.886 | 7.61e-08 | 4 |
| 192.0244 | 0.48 | -0.886 | 7.61e-08 | 4 |
| 427.2059 | 5.22 | -0.879 | 1.34e-07 | 4 |
| 267.0878 | 0.56 | -0.871 | 2.2e-07 | 4 |
| 357.1327 | 0.64 | -0.871 | 2.2e-07 | 3 |
| 414.1695 | 0.63 | -0.871 | 2.2e-07 | 4 |
| 188.0707 | 0.63 | -0.867 | 2.61e-07 | 4 |
| 183.0651 | 0.56 | -0.859 | 4.11e-07 | 3 |
| 216.5644 | 0.76 | -0.859 | 4.11e-07 | 4 |
| 421.2188 | 5.22 | -0.859 | 4.11e-07 | 4 |
| 443.2009 | 5.22 | -0.859 | 4.11e-07 | 4 |
| 275.0766 | 0.77 | -0.855 | 5.36e-07 | 4 |
| 164.0670 | 0.46 | -0.843 | 1.11e-06 | 4 |
| 320.2564 | 7.51 | -0.843 | 1.11e-06 | 3 |
| 623.2420 | 0.63 | -0.839 | 1.37e-06 | 4 |
| 205.0973 | 0.63 | -0.835 | 1.61e-06 | 4 |
| 243.1706 | 5.22 | -0.835 | 1.61e-06 | 4 |
| 216.5645 | 0.84 | -0.824 | 2.95e-06 | 3 |
| 409.1624 | 6.52 | -0.769 | 3.78e-05 | 3 |
| 521.2352 | 5.50 | -0.765 | 4.36e-05 | 4 |
| 191.0539 | 0.75 | -0.761 | 5.08e-05 | 2 |
| 233.5805 | 6.52 | -0.749 | 7.94e-05 | 2 |
| 425.1364 | 6.52 | -0.741 | 0.000106 | 3 |
| 258.2429 | 8.46 | -0.726 | 0.000174 | 3 |
| 438.1256 | 0.67 | -0.726 | 0.000174 | 3 |
| 318.2408 | 8.17 | -0.714 | 0.000255 | 3 |
| 211.5666 | 0.79 | -0.710 | 0.000289 | 4 |
| 170.5406 | 0.83 | -0.706 | 0.000326 | 3 |
| 387.1804 | 6.52 | -0.698 | 0.000408 | 3 |
| 543.2171 | 5.50 | -0.682 | 0.000663 | 2 |
| 180.0867 | 0.47 | -0.679 | 0.000731 | 3 |
| 228.2324 | 10.05 | -0.671 | 0.000903 | 2 |
| 333.5891 | 0.68 | -0.671 | 0.000903 | 3 |
| 708.5122 | 11.89 | -0.667 | 0.000998 | 2 |
| 244.2273 | 7.29 | -0.663 | 0.00109 | 3 |
| 404.2071 | 6.52 | -0.663 | 0.00109 | 3 |
| 254.0937 | 6.52 | -0.651 | 0.00146 | 3 |
| 338.2668 | 6.75 | -0.647 | 0.00158 | 4 |
| 542.6618 | 0.67 | -0.647 | 0.00158 | 4 |
| 273.2619 | 6.26 | -0.639 | 0.00191 | 4 |
| 332.2198 | 6.94 | -0.639 | 0.00191 | 3 |
| 288.2534 | 7.13 | -0.635 | 0.0021 | 4 |
| 272.2585 | 6.26 | -0.631 | 0.0023 | 4 |
| 254.2477 | 6.26 | -0.628 | 0.00247 | 4 |

| m/z | RT (min) | rho | FDR | Steps |
|---|---|---|---|---|
| 469.7800 | 5.24 | -0.628 | 0.00247 | 3 |
| 263.1156 | 5.22 | -0.620 | 0.00292 | 2 |
| 271.1653 | 6.03 | -0.616 | 0.00318 | 3 |
| 432.2385 | 6.52 | -0.612 | 0.00344 | 2 |
| 527.2222 | 5.52 | -0.600 | 0.00449 | 2 |
| 220.1153 | 5.38 | -0.596 | 0.00487 | 2 |
| 282.2792 | 11.20 | -0.588 | 0.00566 | 3 |
| 437.1940 | 7.01 | -0.588 | 0.00566 | 2 |
| 271.1130 | 5.22 | -0.584 | 0.00597 | 2 |
| 320.2564 | 8.87 | -0.584 | 0.00597 | 3 |
| 261.1098 | 6.09 | -0.577 | 0.00685 | 2 |
| 194.1154 | 5.19 | -0.545 | 0.0125 | 3 |
| 288.2534 | 8.17 | -0.541 | 0.0133 | 3 |
| 381.7269 | 1.39 | -0.541 | 0.0133 | 3 |
| 300.1870 | 6.33 | -0.537 | 0.014 | 3 |
| 436.7757 | 5.16 | -0.537 | 0.014 | 3 |
| 444.3325 | 8.52 | -0.537 | 0.014 | 3 |
| 241.1548 | 5.80 | -0.533 | 0.0149 | 3 |
| 170.5405 | 0.67 | -0.526 | 0.0171 | 3 |
| 384.3083 | 6.20 | -0.514 | 0.0209 | 3 |
| 216.5997 | 6.03 | -0.506 | 0.0236 | 3 |
| 534.6729 | 0.64 | -0.502 | 0.025 | 3 |
| 215.0981 | 8.80 | -0.490 | 0.0299 | 2 |
| 298.2743 | 8.74 | -0.490 | 0.0299 | 3 |
| 683.2879 | 5.36 | -0.490 | 0.0299 | 4 |
| 244.2635 | 6.32 | -0.482 | 0.0335 | 2 |
| 304.2847 | 6.37 | -0.475 | 0.0372 | 3 |
| 425.7533 | 5.06 | -0.467 | 0.0415 | 3 |
| 218.2117 | 4.91 | -0.459 | 0.0462 | 3 |

Of the 161 features, 144 (89.4%) changed in the expected direction in at least three of four consecutive dose steps and 61 (37.9%) were strictly monotonic across all concentrations; no feature showed fewer than two consistent steps.

### Supplementary Table 7 | Complete targeted metabolomic profiling of B. subtilis biofilm specialised metabolites

| Metabolite | *m/z* | Adduct | Mode | Norm. | FC | S vs C | S vs Q | Q vs C |
|---|---|---|---|---|---|---|---|---|
| **Bacillaene** | 581.358 | $[M+H]^+$ | ESI+ | Terfenadine | **1.67×↑** | **0.0042**** | **0.0084**** | 0.47 |
| Surfactin C13 | 1008.659 | $[M+H]^+$ | ESI+ | Terfenadine | 1.28×↑ | 0.27 | 0.61 | 0.96 |
| **Surfactin C14** | 1022.670 | $[M+H]^+$ | ESI+ | Terfenadine | **1.67×↑** | **0.020*** | 0.32 | 0.49 |
| Surfactin C15 | 1036.690 | $[M+H]^+$ | ESI+ | Terfenadine | 1.37×↑ | 0.21 | 0.49 | 0.96 |
| **Bacillaene** | 579.344 | $[M-H]^-$ | ESI− | Chlorpropamide | **1.59×↑** | **0.0069**** | **0.0030**** | 0.97 |
| Surfactin C13 | 1006.645 | $[M-H]^-$ | ESI− | Chlorpropamide | 1.97×↑ | 0.16 | 0.074 | 0.91 |
| Surfactin C14 | 1020.661 | $[M-H]^-$ | ESI− | Chlorpropamide | 1.54×↑ | 0.37 | 0.089 | 0.53 |
| Surfactin C15 | 1034.676 | $[M-H]^-$ | ESI− | Chlorpropamide | 1.46×↑ | 0.45 | 0.15 | 0.72 |
| **Plipastatin C14** | 730.390 | $[M-2H]^{2-}$ | ESI− | Chlorpropamide | **1.31×↑** | **0.044*** | **0.037*** | 0.64 |
| Plipastatin C15 | 737.400 | $[M-2H]^{2-}$ | ESI− | Chlorpropamide | 1.30×↑ | 0.085 | 0.086 | 0.75 |
| Plipastatin C16 | 744.410 | $[M-2H]^{2-}$ | ESI− | Chlorpropamide | 1.13×↑ | 0.34 | 0.19 | 0.53 |
| Plipastatin C17 | 751.414 | $[M-2H]^{2-}$ | ESI− | Chlorpropamide | 1.12×↑ | 0.44 | 0.24 | 0.60 |
| Bacillibactin | 881.248 | $[M-H]^-$ | ESI− | Chlorpropamide | 0.89×↓ | 0.77 | 0.99 | 0.68 |

All metabolites measured by LC-HRMS in both positive (ESI+) and negative (ESI−) ionization modes. FC, fold change (SAIRGA/Control, linear scale). All pairwise P values are Games–Howell-adjusted, following Welch's ANOVA. S, SAIRGA; Q, SAIRGQ; C, untreated control. n = 6 biological replicates per group. Positive-mode measurements were normalized to terfenadine (CV 0.34–0.60%, one-way ANOVA P = 0.53) and negative-mode measurements to chlorpropamide (CV 10.7%, one-way ANOVA P = 0.16). Bacillaene and the surfactin C13–C15 congeners were quantified in both ionization modes; Fig. 5e plots one measurement per compound, selected by within-group precision, so the three negative-mode surfactin rows are reported here but not plotted. Metabolites reaching significance are shown in bold. *P < 0.05; **P < 0.01. Light-grey P values denote non-significant comparisons. Thin grey lines separate ionization modes and metabolite classes.
